# Multiscale dynamics underlying neocortical slow oscillations

**DOI:** 10.1101/2021.06.09.447804

**Authors:** Maurizio Mattia, Maria Perez-Zabalza, Núria Tort-Colet, Miguel Dasilva, Alberto Muñoz, Maria V. Sanchez-Vives

## Abstract

Slow oscillations in the sleeping and anesthetized brain invariantly emerge as an alternation between Up (high firing) and Down (almost quiescent) states. In cortex, they occur simultaneously in cell assemblies in different layers and propagate as traveling waves, a concerted activity at multiple scales whose interplay and role is still under debate. Slow oscillations have been reported to start in deep layers, more specifically in layer 5. Here, we studied the laminar organization of slow oscillations in the anesthetized rat cortex and we found that the activity leading to Up states actually initiates in layer 6, then spreads towards upper layers. Layer 5 cell assemblies have a threshold-like activation that can persist after layer 6 inactivation, giving rise to hysteresis loops like in “flip-flop” computational units. We found that such hysteresis is finely tuned by the columnar circuitry depending on the recent history of the local ongoing activity. Furthermore, thalamic inactivation reduced infragranular excitability without affecting the columnar activation pattern. We propose a role for layer 6 acting as a hub unraveling a hierarchy of cortical loops.

## INTRODUCTION

Primary sensory cortices like visual cortex (V1), display powerful computational capabilities. A prominent example is stimulus selectivity, which relies on intrinsic nonlinear properties of cortical neurons and on feed-forward thalamocortical circuits^1,2^. Nonlinear network dynamics also underlie the ability of V1 to sustain input-related activity once visual stimuli are removed^3^ or when the stimulation sequence is incomplete^4^, a computational primitive allowing integration of information across time. These intrinsic nonlinear dynamical features are expressed both during sensory processing and under resting conditions. In part this is due to the fact that both spontaneous and evoked spatiotemporal activity patterns reflect the same underlying structure. For instance, orientation maps spontaneously pop up ordered in time in V1 of anesthetized cats^5^, and during the Up high-firing states of slow oscillations (SO), activity patterns similar to stimulus-evoked responses are randomly replayed both in auditory and somatosensory cortices of anesthetized rats^6^. This is the result of a collective nonlinear dynamics shaped by intra-cortical connectivity, which does not change across different brain states like wakefulness and slow-wave activity under deep anesthesia and non-REM sleep^7^. Understanding the mechanistic origins of such computational primitives is a challenge of central interest and SO are an open window to explore them.

The multiscale nature of SO, which simultaneously involves several interacting dynamical systems, adds up to the challenge^8^. At the micro- and mesoscopic level of *in vivo* neurons and localized neuronal assemblies, respectively, SO occur as an alternation of relatively high firing rate periods (Up states) and almost quiescent ones (Down states)^9–12^. These are not isolated dynamical patterns: at the macroscale, Up and Down states propagate as slow traveling waves along the cortical surface determining an effective synchronization between SO of nearby local networks^7,11,13,14^. Besides, SO are found in the thalamus as well, with state onsets locked in time with the cortical ones^15^. Although SO are widely considered to be originated in neocortex, given that the cortical tissue alone can generate them^16,17^ while isolated thalamus cannot^18^, it is still widely debated how the thalamus actively affects slow-wave activity^8,12,15,19^.

An additional level of complexity is added if we consider the cortical column, where SO are differently expressed across cortical layers^10,16,20^. Neurons in layer 5 (L5) are the first to prime Up state onsets, and display the highest firing rates in cortical slices^16^, similar to what has also been found *in vivo*^10,20^. Furthermore, optogenetic or local stimulation of L5 neurons elicits Up states in an all-or-none manner similar to those spontaneously occurring during SO^13,16,21,22^. This is not the case of supragranular neurons in layer 2/3 (L2/3) when similarly activated^21,22^, highlighting a vicarious role of superficial layers. This is also compatible with the evidence of an upward spreading of Up state activation from deep layers to L2/3^10,16,20^. This is why L5 assemblies are thought to initiate and drive SO, although the underlying nonlinear dynamics have not yet been fully characterized^23–25^.

In recent years, deep infragranular layer 6 (L6) has been revealed to have a critical role in integrating multiscale information coming from the thalamus and both lateral and top-down intra-cortical connectivity^26–33^. In spite of such role in cortical processing, L6 activation during SO seems to occur always after Up state onset in L5^16,20,34^. All these evidence opens questions regarding the interplay between L5 and L6, a possible paradox to unravel.

Here we carried out a detailed quantitative exploration of the multiscale dynamics of SO to understand the rules of columnar function. After confirming the leading role of L5 neuronal pools in generating and sustaining Up states, we demonstrate a history-dependent attractor dynamics likely mediated by synaptic reverberation and activity-dependent adaptation mechanisms. We find evidence that L6 provides a critical input to L5 to elicit the transition from Down to Up state. Our findings reconcile the proposed role of L6 as an integrator of synaptic input from thalamus and other cortical areas and the intra-columnar information flow with the intrinsic nonlinear properties of the columnar circuit. Finally, to identify whether the role that we attribute to L6 has a main thalamic component, we inactivate the connected thalamus, demonstrating that the inter-laminar dynamics are mainly driven by the cortical inputs, although the thalamus has a modulatory role.

## RESULTS

We recorded the synaptic and neuronal activity underlying slow Up/Down oscillations (SO) from the visual cortex (V1) of anesthetized rats (n = 18 adult male Wistar rats, oscillation frequency 1.11 ± 0.16 Hz, mean ± s.d.). To this end we collected unfiltered raw signals using a 16-channel silicon probe whose position with respect to the laminar distribution was histologically identified in an additional set of experiments (**Fig. 1a**, n = 5 adult male Wistar rats and Online Methods). From each channel of the silicon probes we extracted low-pass filtered local field potentials (LFPs) and multi-unit activities (MUAs) across all cortical layers (**Fig. 1b**, and Online Methods).

**Figure 1.**
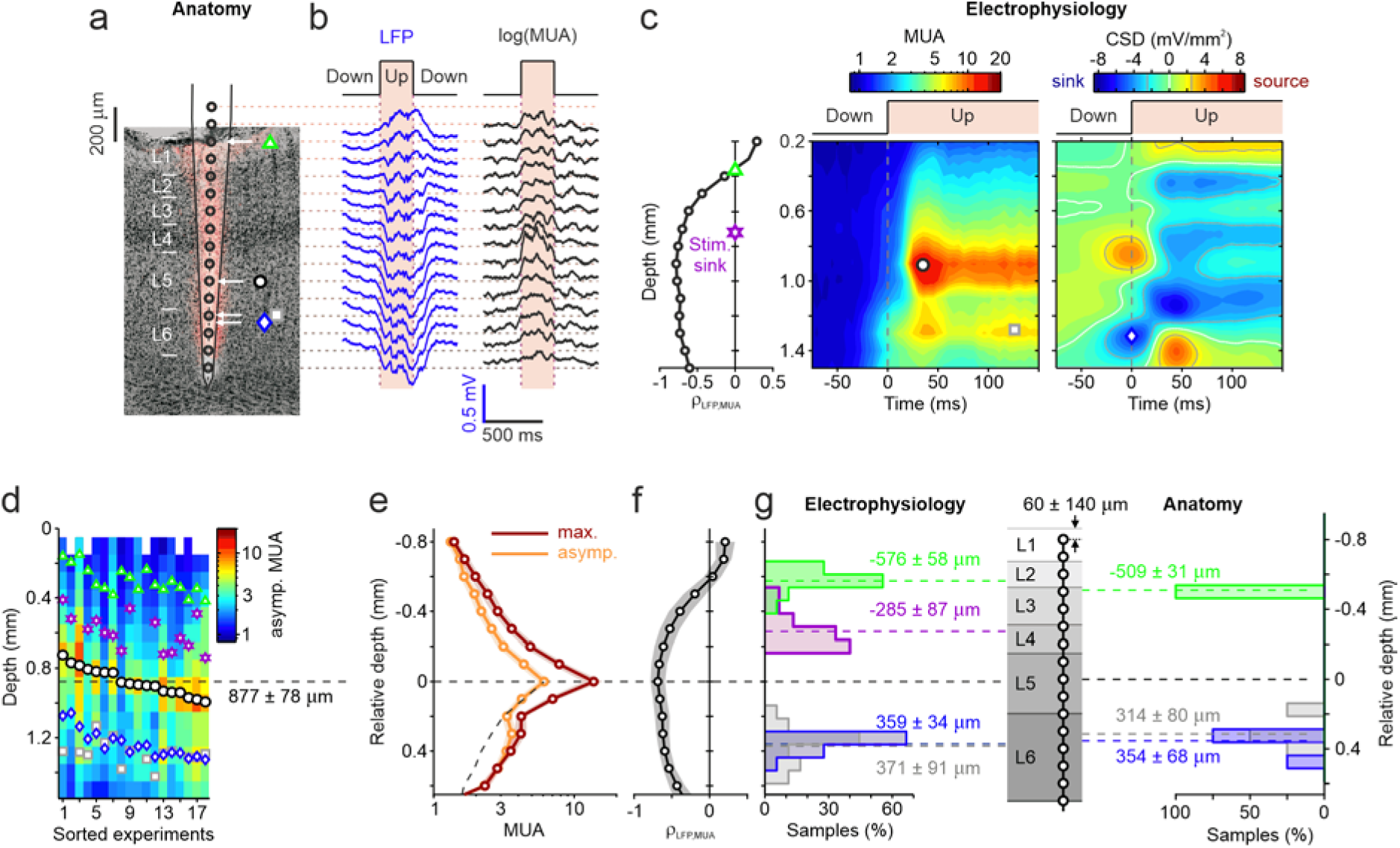
Electrophysiological markers of SO across cortical layers. (**a**) 16-channel silicon probe (inter-electrode distance: 100 μm) and histological identification of the V1 cortical layers of anesthetized rats from an example experiment with simultaneous anatomical and electrophysiological characterization (n = 5). Symbols as in **c**. (**b**) Example LFPs and MUAs extracted at different cortical depths in V1. Unfiltered raw signals were recorded from each channel. (**c**) Left, correlation coefficient ρ between LFP and log(MUA) at different cortical depths. Green triangle, depth where LFP polarity reversal occurred. Purple star, depth of L4 estimated from CSD evoked by photic stimulation (Online Methods). Center, average MUA centered around detected Down-to-Up transitions. Black circle, time and depths where maximum MUA was found. Gray square, depth of the second MUA peak during Up states. Right, average CSD centered around upward transitions. Red, sources; blue, sinks. Blue diamond, depth of the CSD sink preceding Down-to-Up transitions. (**d**) Average asymptotic MUA for all recordings (n = 18). Asymptotic MUA was the average activity during Up states without the first 100 ms. First electrode (depth = 0 μm) was aligned to the cortical surface. Symbols as in **c**. Experiments sorted by depths of maximum MUA peaks. (**e**) Average across experiments of maximum (red) and asymptotic (orange) MUA *versus* relative depth. Shadings, s.e.m. Dashed line, mirrored asymptotic MUA at positive relative depths. Black curve, distance between black dashed and orange curves. (**f**) Average across experiments of the correlation ρ between LFP and log(MUA). Gray strip, s.d. (**g**) Left, relative depth distributions of electrophysiological markers in **d** (n = 18). Center, layer depths estimated as mean across experiments with anatomical characterization (n = 5) together with the average displacement of the silicon probe based on the distance of the most superficial channel from the cortical surface (60 ± 140 *μ*m). Right, relative depth distributions of the electrophysiological markers identified by histology (n = 5). Relative depths in panels (**e**-**g**) were the distances from corresponding maximum MUA positions. Average depths, mean ± s.d.

### The fingerprints of SO across cortical layers

For our purposes it was critical to assure the location of the electrodes across the six cortical layers. By using different measures (for details see Online Methods), we obtained a reliable depth information of several electrophysiological markers that resulted in a register of depth distributions (**Fig. 1g**). Besides manual alignment of the first electrode in the arrays to the cortical surface, we firstly measured the depth of the LFP polarity reversal, known to occur between 0.25 and 0.5 mm^10,35^, and thus pointing out the supragranular layer 2/3 (L2/3). This depth corresponded to the sign change in cross-correlation between LFP and MUA (**Fig. 1c**, left). Moreover, we resorted to a current source density (CSD) analysis^36^ of stimulus-evoked responses to detect the current sink (purple star in **Fig. 1c**, left) corresponding to the lateral geniculate nucleus (LGN) projections to layer 4 (L4)^37^ (Online Methods). During SO, two more depth markers were apparent in the onset-triggered average of MUA (**Fig. 1c**, center): i) the maximum MUA transiently occurring at the beginning of the Up state and ii) a deeper second peak of activity reliably detectable from the asymptotic MUA after 100 ms from the Up state onset. With the exception of this second deeper peak of MUA, all these markers reliably replicate what recently found in naturally sleeping mice^38^. Finally, from the SO onset-triggered average of the CSD, a deep sink with its coupled more superficial source systematically appeared before the Down-to-Up transitions at 1,300 μm and 850 μm depth respectively (**Fig. 1c**, right). All these features were found in the large majority of the performed experiments (**Fig. 1d**). Remarkably, the depth markers were at an almost constant distance from the maximum MUA depth found at an average depth of 877 ± 78 μm (mean ± s.d.). Using such depths in each experiment as reference, the inter-experiment variability was relatively small both in the asymptotic and maximum MUA (**Fig. 1e**), and in the LFP-MUA correlation (**Fig. 1f**). With this realignment (**Fig. 1g**, left), the depth marker from the stimulus-evoked CSD remarkably pointed out L4, as expected, while the LFP polarity reversal always occurred in L2/3 once the laminar distribution estimated from the histological measures was adopted (**Fig. 1g**, center). The maximum MUA was consistently found in L5 as in refs.^10,16,38^, while the both unexpected early active deep CSD sink and second MUA peak were found at the same depth within L6. These localized distributions of marker depths in different cortical layers was eventually confirmed by a set of additional experiments (n = 5) where electrophysiological measures were co-registered with histological localization of both the silicon probes and the layer depths in the V1 of rats (**Fig. 1g**, right).

### Ordered Up state activation from deep to superficial layers

The presence of a synaptic activation associated to the CSD sink in L6 consistently preceding a L5 MUA activation (**Fig. 1c**, right) supports the hypothesis of a trigger for the Up state initiation outside the boundary of L5, contradicting what has been previously reported in anesthetized mammals^10,20^ and in acute cortical slices^16,22,39^. We further investigated such possibility by directly inspecting the patterns of activation times of MUA across cortical layers. For each detected Up state, we extracted an array of time lags between the activation times in all electrodes and the MUA onset times with respect to L5, which was considered zero lag. On the resulting matrix with all Down-to-Up transitions, a principal component (PC) analysis was performed, allowing to sort the time-lag matrix with respect to the first PC (**Fig. 2a**, and Online Methods). Across all the experiments, we consistently found the same picture: a gradient of negative time lags in deeper electrodes corresponding to L6, meaning that the firing in L6 neurons preceded that in L5 up to a maximum of 45 ms. By pooling the sorted Down-to-Up transitions (438 ± 85; mean ± s.d.; n = 18 experiments) in five equally-sized groups, average MUA and CSD highlighted the chain of events in time characterizing the Down-to-Up transition across a cortical column (**Fig. 2b**). Firstly, a CSD sink appeared in L6 together with its coupled source. Almost simultaneously a MUA increase started at the same depth. Only after a variable delay of several milliseconds a sudden upward transition occurred also in L5, from which followed the spread of activity towards the more superficial layers, L4 and L2/3.

**Figure 2.**
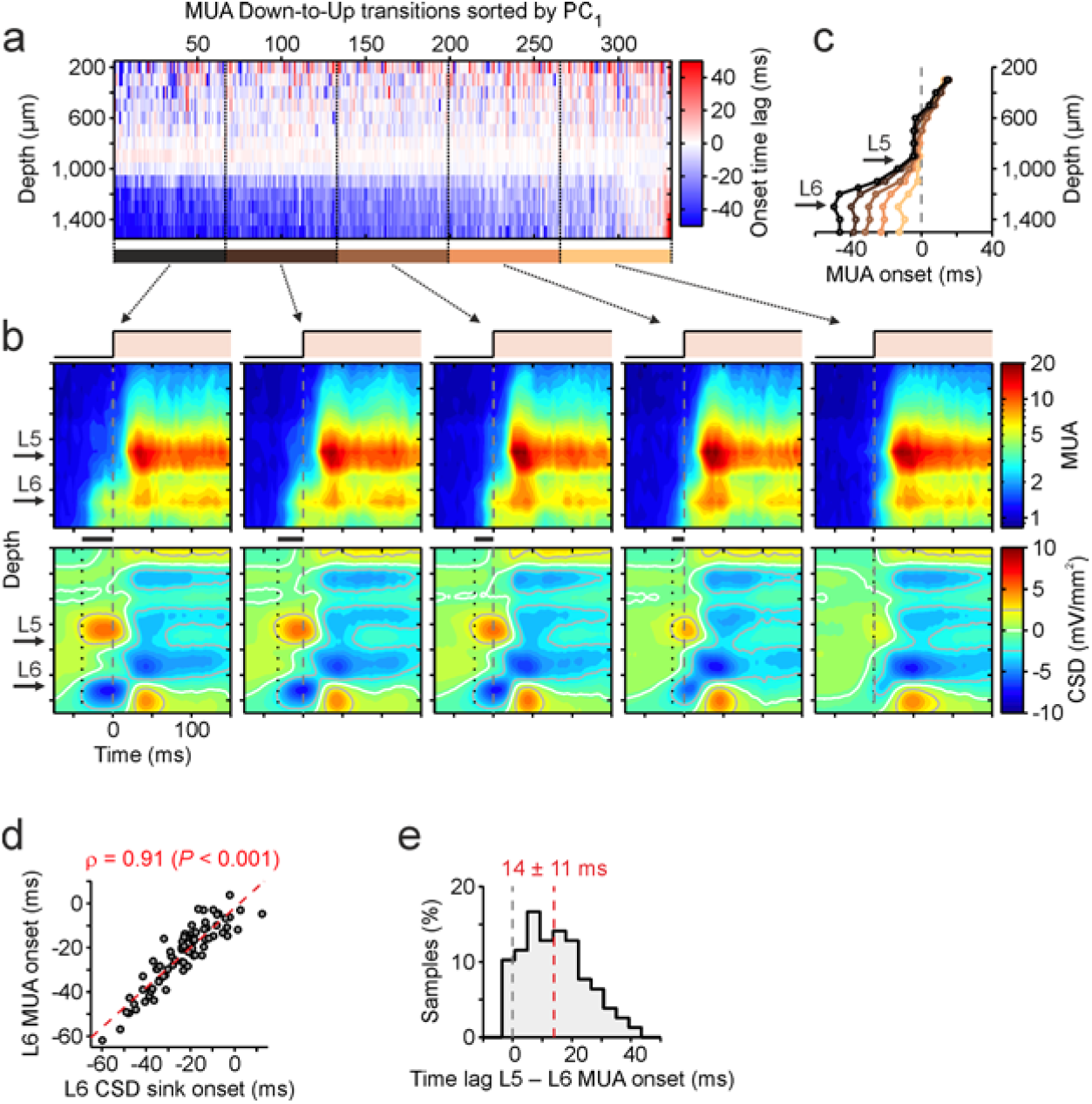
Synaptic and firing activity onset across lamina. (**a**) MUA onset time lags across cortical depth of the same example recording as in **Figure 1c**. Each column of the matrix was a detected Down-to-Up transition and time lags were from average onset times over the three electrodes around maximum MUA depth (L5). Upward transitions were sorted with respect to the first component from a PCA on the time lag matrix. (**b**) Sorted upward transitions pooled in 5 groups from which average MUA (top) and CSD (bottom) were carried out around Up state onset in L5 (t = 0 ms). Black horizontal bars, time lags between L6 CSD sink onset (vertical dotted line) and L5 MUA onset times. (**c**) Average MUA onset time over grouped upward transitions for the same example recording. Down-to-Up transitions are now sorted by the time lag between L6 and L5 MUA Up state onsets. (**d**) Correlation between L6 MUA and CSD sink onset times of grouped Up states from all experiments (n = 78 transition groups). (**e**) Histogram of time lags between L6 and L5 MUA onset times from all grouped transitions. Average time lag (red dashed line), mean ± s.d.

Such spatiotemporal pattern in the cortical column activation was even more obvious when looking at the profiles of the average time lags between MUA crossings of a fixed low threshold, and the previously detected L5 MUA activation (**Fig. 2c**, and Online Methods). To further reduce the variability of the measures, here the five Up state groups were pooled from MUA upward transitions sorted directly by L6-to-L5 MUA onset delays. The activation spread from L5 to L2/3 was fast (average 18 ms) and rather constant, while the former MUA onset in L6 displayed a widely variable time lag (**Fig. 2c**). The wide distribution of activation times in L6 and the simultaneous onset of synaptic input and MUA at the same depth were found also at population level (**Fig. 2d**). L5 MUA onset consistently followed the activation time in L6 in all Up state groups from all experiments (**Fig. 2e**), with an average lag of 14 ± 11 ms (mean ± s.d.).

### Evidence of L5 persistent activity around Up state offsets

The end of the Up state and beginning of the Down state was characterized by the end of the firing (**Fig. 3a-b**). While the Down-to-Up transition was led by infragranular layers, the transition from Up to Down state started rather in superficial layers spreading from there towards L5 and L6 (**Fig. 3c**). Both L5 and L6 shared the role in terminating the Up-to-Down transition, leading the end of the firing at 50% (**Fig. 3d**, both positive and negative time lags).

**Figure 3.**
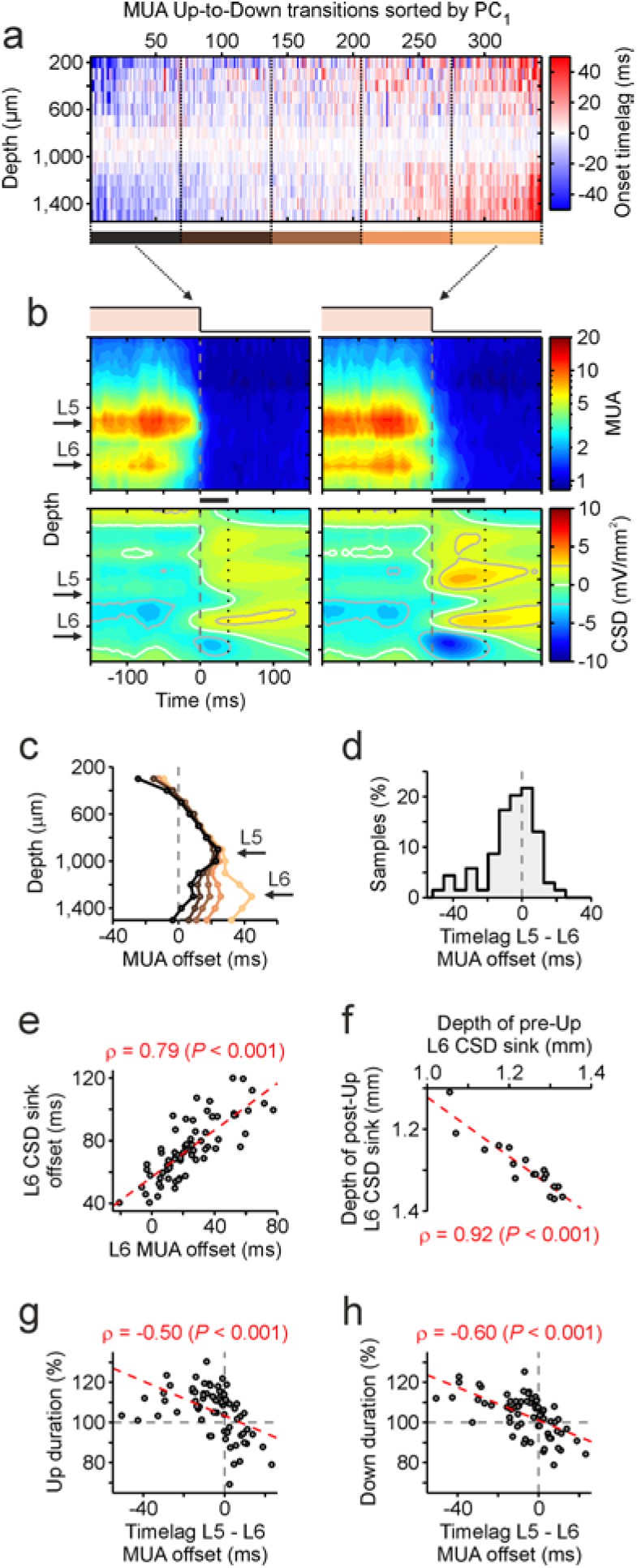
Synaptic and firing offset during Up state termination. (**a**) MUA offset time lags across cortical depth of an example recording, the same as in **Figure 1** and **2**. Columns of the time lag matrix are the detected Up-to-Down-transitions. PCA-based sorting and time lags from L5 MUA offset as in **Figure 2a**. (**b**) Average MUA (top) and CSD (bottom) of the first (left) and last (right) group of sorted downward transitions. Horizontal bars, time lags between L5 MUA inactivation times and L6 CSD sink offset (vertical dotted line). (**c**) Average MUA offset time of the above 5 grouped transitions, as in **Figure 2c**. (**d**) Histograms of L6-to-L5 MUA offset delays, as in **Figure 2e** (n = 69 transition groups). (**e**) Correlation between MUA and CSD offset times in L6, across grouped transitions. (**f**) Depths of the CSD sink in L6 before and after Up states, from all experiments (n = 18). (**g**-**h**) L6-to-L5 MUA offset delay *versus* average Up (**g**) and next Down (**h**) state duration for each transition group. Gray dashed lines, axes origins.

When looking into the CSD, the laminar synaptic activity around the Up state offset also displayed characteristic spatiotemporal patterns: deep CSD sinks emerged at the beginning of Down states (**Fig. 3b**, bottom), lasting on average for 51 ± 12 ms (mean ± s.d.) after MUA offset in L6. Both, L6 CSD and MUA offset showed a tight correlation (ρ = 0.79, P < 0.001) at the level of the downward transition groups (**Fig. 3e**, n = 69), as it did for Down-to-Up transitions (**Fig. 2d**). CSD sinks in L6 preceding and following Up states were found at the same cortical depth in all recordings (**Fig. 3f**, n = 18), suggesting that both L6 CSD sinks could be fingerprints of the same input synaptic activity.

This view is compatible with another evidence collected across all experiments, the anti-correlation (*ρ* = −0.50, *P*< 0.001) between the duration of Up states and the time lag between Up state offset in L5 and L6 (**Fig. 3g**). In other words, for short Up states the firing rate in L6 finished before that in L5. However, for longer Up states, the firing rate in L6 persisted after the end of firing in L5. Therefore, in the termination of the Up states, there are two possibilities regarding the L5-L6 lag, positive or negative. However, at the initiation of the Up state or Down-to-Up transition, L6-to-L5 onset delays were only positive (**Fig. 2e**).

L6-to-L5 offset delays at the termination of the Up state were also tightly anti-correlated (*ρ* = −0.60, *P*< 0.001) with the average duration of the subsequent Down state (**Fig. 3h**). In particular, longer Up states, that corresponded to an earlier L5 termination with a longer persistence of L6 activity, were followed by longer Down states. An activity-dependent fatigue mechanism such as hyperpolarizing K^+^ currents underlying Down states^16,40^ could support such relationship.

### Hysteresis loop in L5 cortical modules

The systematic anticipation of L6 MUA onset with respect to MUA upward transitions in L5, together with the early appearance of a synaptic input in L6 at the end of Down states (**Fig. 2**), were compatible with a feed-forward synaptic connection from L6 to L5 cortical modules which is supported by anatomical studies^26,41^. In this framework, the availability of simultaneous L5 MUA and its potential input, L6 MUA, allowed us to inspect the input-output relationship of L5 cortical modules during SO. We focused in this section on short Up states (20% of briefest durations), where L5 high MUA persisted after L6 offset, a possible footprint of state-dependent nonlinear dynamics proposed for information processing in sensory areas^42^. Nonlinear dynamics was confirmed by the evidence of a hysteresis loop (**Fig. 4a**, bottom-left), such that in response to the same input (L6 MUA) L5 generated different outputs depending on the phase of the Up/Down cycle. Intriguingly both Down-to-Up and Up-to-Down (right and left steep segments of the loop, respectively) seemed to occur once a threshold value of the input activity was reached, bringing to a sudden change in the trajectory in L6-L5 MUA plane. The two thresholds were shifted, such that higher firing rate of L6 was needed to initiate an Up state in L5 than to bring back L5 from Up to Down state, highlighting the history-dependence of L5 module dynamics.

**Figure 4.**
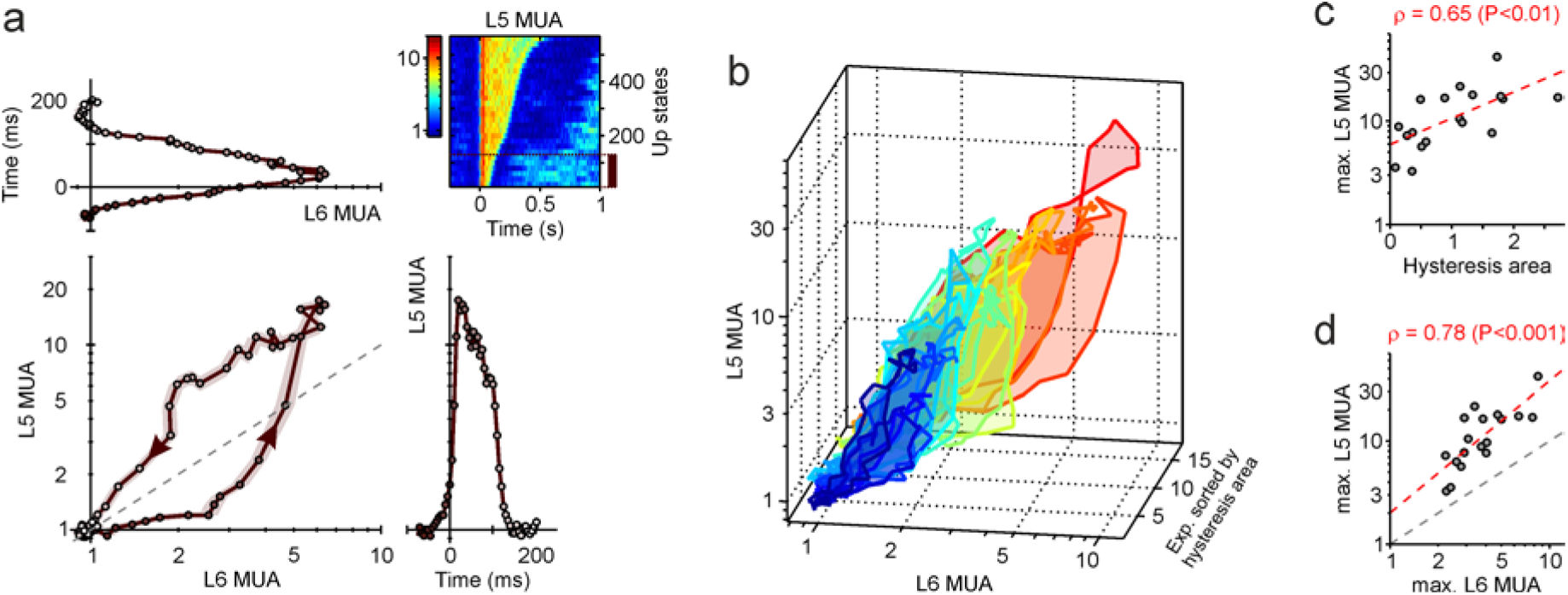
Hysteresis loop of L5 cortical modules in response to Up/Down oscillations in L6. (**a**) Up state onset-centered average MUA of two electrodes at L6 and L5 depth (top-left and bottom-right panels) of an example recording. Circles are the samples collected every 5 ms, with shadings from dark brown to white representing increasing time course. Average MUAs are from the first of five groups of Down-to-Up transitions detected in L5 with the shortest next Up state duration (top-right panel). MUA profiles are obtained concatenating average MUA centered around Down-to-Up and Up-to-Down transitions and for each of them considering only a time period half of the average Up state duration in the selected group (Online Methods). (**b**) Average hysteresis loop as in **a** for all the n = 18 experiments, sorted by ascending hysteresis area (color codes for ranking). (**c**) Correlation between maximum MUA in L5 and hysteresis areas across all experiments. (**d**) Maximum MUA in L6 and L5 for all experiments. Red dashed line, linear regressions as in **c**.

The features highlighted for the example recording were found also in the majority of the experiments, specifically those with a large enough area inside the hysteresis loop (**Fig. 4b**). In particular, an exponential relationship was apparent between hysteresis area and the maximum firing rate in L5 during Up state (**Fig. 4c**, ρ = 0.65, *P* < 0.01): the larger the Up state firing rate was, the larger the hysteresis loop, and hence more distant were the thresholds L6 MUA needed to elicit upward and downward transitions in L5. This observation further supported the hypothesis that hysteresis in L5 resulted from nonlinear dynamics: the larger was the capability to separate Up and Down state activity, and hence for L6 MUA to amplify Up/Down oscillations, the less susceptible were L5 cortical modules to change their current state. As well, the excitability/nonlinearity of L5 cortical modules resulted to be related to the maximum MUA in L6 (**Fig. 4d**, ρ = 0.78, *P* < 0.001) with a power law *r*_L5_ = 2.0 *r*_L6_^1.3^ (with *r*_L*x*_ being the MUA in layer *x*).

### History-dependent attractor dynamics behind L5 hysteresis

We hypothesized a local network dynamics where Up and Down states were the preferred and attracting activity conditions based on the L6-L5 hysteresis. In order to investigate such mechanistic scenario, we resorted to an effective minimal rate model embodying the experimental features of infragranular cortical layers. To this purpose, L6 and L5 of a cortical column were described as two separated pools of neurons with the L6 feeding its activity as excitatory synaptic input to L5 (**Fig. 5a**). We described the L6 module as a “relay” station, mapping almost linearly the synaptic input received from other cortical areas or thalamic inputs into the local MUA.

**Figure 5.**
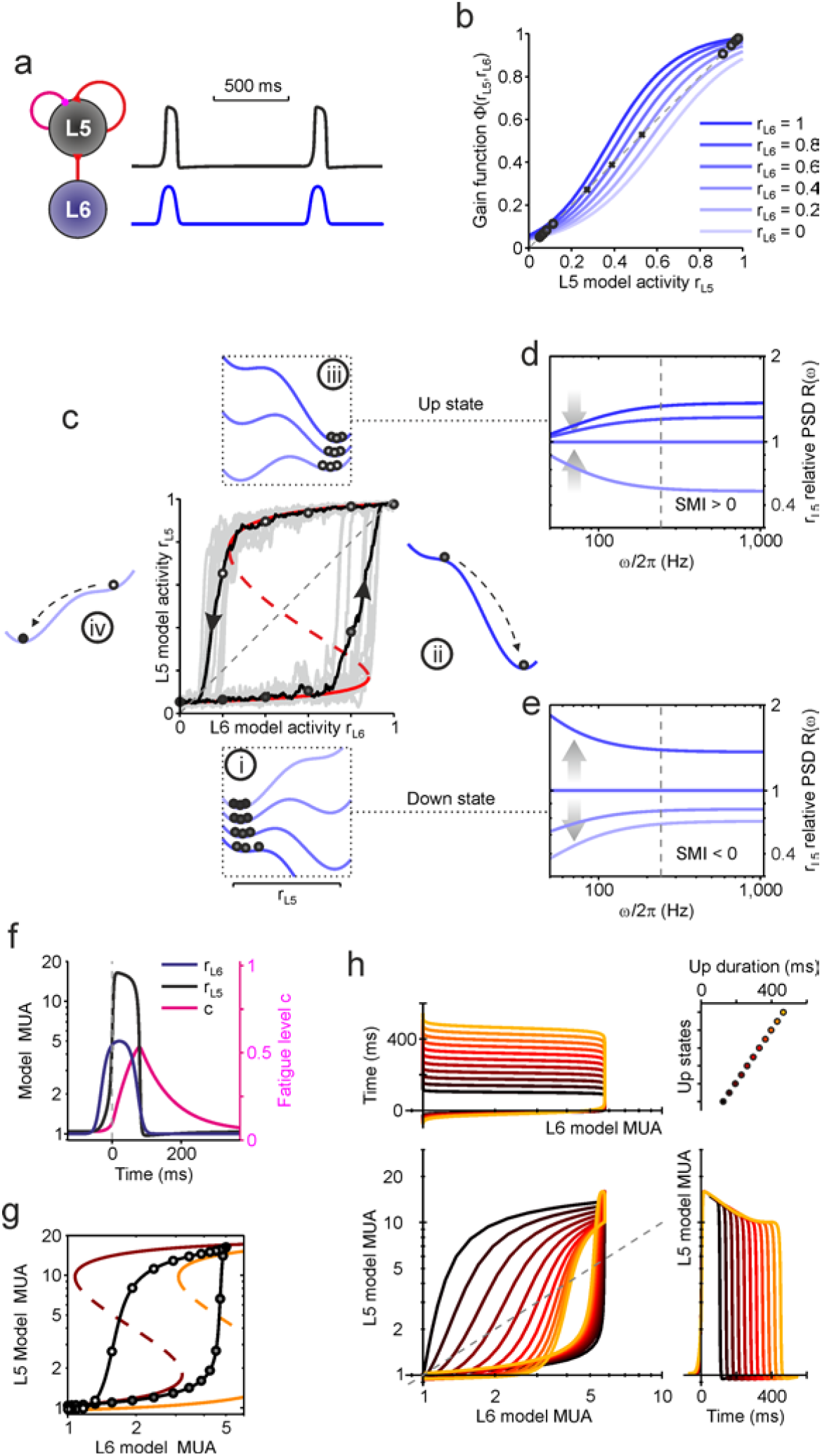
History-dependent nonlinear dynamics in a L6-L5 minimal model. (**a**) Minimal rate model composed of a L6 module projecting its activity *r*_L6_ through excitatory synapses to a L5 module with gain function Φ shaped by recurrent excitatory synapses (red arrow). Right, example traces of L6 (blue) and L5 (black) firing rates. Purple arrow, activity-dependent self-inhibition modeling spike frequency adaptation. (**b**) Sigmoidal Φ of L5 module at different *r*_L6_ values. Dashed line, input equals output rates. Circles (crosses), stable (unstable) fixed point of *r*_L5_ dynamics. (**c**) Example trajectories in the (*r*_L6_,*r*_L5_) plane showing hysteresis loops (gray traces). Firing rates fluctuate due to finite-size noise. Red curve, nullcline d*r*_L5_/dt = 0 of stable (solid) and unstable (dashed) fixed points. Black track, average trajectory. Subpanels **i**-**iv**, changes of L5 effective energy landscape. Circles representing L5 activity. Blue shadings as in **b**. (**d**-**e**) Relative power spectral density (PSD) *R*(ω), theoretically estimated at different *r*_L6_. Reference PSD is at *r*_L6_ = 0.6. (**d**) L5 during Up state, loop phase **iii**. (**e**) L5 during Down state, loop phase **i**. Dashed lines separate high- and low-frequency bands used to compute the spectral modulation index SMI. (**f**) L5 onset-centered *r*_L5_, *r*_L6_ (left axis) and fatigue variable *c* modulating L5 activity-dependent self-inhibition (right axis, purple as in **a**). (**g**) Example trajectory for the activity plotted in **f**. Dark red (orange) curve, nullcline at the Up state onset (offset), when *c* is minimum (maximum) in **f**. (**h**) Model activity *r*_L5_ (bottom right) and *r*_L6_ (top left) for different Up state durations (top right), and the resulting trajectories in the (*r*_L6_,*r*_L5_) plane (bottom left).

We modeled L6 activity *r*_L6_ as periodically oscillating between high- and low-firing states with residence periods comparable to those measured from experiments (Online Methods). In turn, L5 module was modeled to have firing rate *r*_L5_(*t*) with a first-order dynamics driven by incoming synaptic current via a gain function Φ: τ d*r*_L5_/dt = Φ(*r*_L5_, *r*_L6_) − *r*_L5_. The contribution to L5 synaptic current were a feed-forward input from L6 module and its own activity *r*_L5_ through recurrent synaptic connections. Strong excitatory synaptic reverberation can yield to sigmoid-shaped Φ (**Fig. 5b**), allowing the coexistence of two attractor states^43^. In other words, in both states L5 firing rate *r*_L5_ self-consistently reproduced itself and any small perturbation was dampened [Φ(*r*_L5_, *r*_L6_) = *r*_L5_ and Φ′ = ∂Φ/∂ *r*_L5_ < 1, respectively; **Fig. 5b**, circles]. In this framework, L6 activity could be seen as a modulator of L5 excitability: the higher *r*_L6_ was, the more excitable L5, which roughly corresponded to a Φ more shifted to the left in **Figure 5b** and thus to attractor states at relatively higher frequencies. We aimed to model finite-size cell assemblies, thus with endogenous fluctuations of firing rate *r*_L5_. These fluctuations were embodied in *r*_L5_ dynamics by considering its finite-size counterpart *r*_L5_^(N)^ = *r*_L5_ + *W*(*t*), which in turn determined the recurrent synaptic input, changing the driving gain function into Φ(*r*_L5_^(*N*)^, *r*_L6_) (Online Methods). *W*(*t*) was a white noise with zero mean and variance *r*_L5_/*N*, and *N* was the number of neurons in L5 modules as in ref. ^44^, such that larger was the network size *N*, smaller was the finite-size fluctuations of *r*_L5_^(*N*)^.

Under the influence of the oscillatory input provided by the L6 module, the above minimal rate model of L5 was capable to display a hysteresis loop resembling those measured *in vivo* (**Fig. 5c**). Irrespective to finite-size fluctuations of single Up/Down cycles (gray traces), the average trajectory (black) showed four distinct phases (**Fig. 5c i-iv**), as in (**Fig 4a**). These stages can be described following the changes in the effective energy landscape on top of which L5 rate model wander. The effective energy (Lyapunov function) was defined as the integral of the force field Φ(*r*_L5_, *r*_L6_) − *r*_L5_ driving the changes in time of *r*_L5_^44,45^. In response to the rise of L6 activity, *r*_L5_ only mildly changed because the system was trapped in the Down attractor state (**Fig. 5c i**). Further increasing *r*_L6_ made the valley shallower and the state less stable, until valley bottom (solid red curve) disappeared making the Up valley the only available state rapidly reached by the system (**Fig. 5c ii**). Once L6 activity decreased to come back to its Down state, the Up attractor valley was in turn made less stable, shrinking the distance to the ridge (dashed red curve) of the barrier between the two valleys and with only a slight change of *r*_L5_ (**Fig. 5c iii**). Last phase of the loop occurred when *r*_L6_ was small enough to have only one stable attractor, the Down state, where the system suddenly dropped (**Fig. 5c iv**).

In this theoretical framework, L6 activity modulated the bistability of L5 cortical modules (i.e. the simultaneous coexistence of Up and Down states), by shaping its effective energy landscape. Following what is observed in the *in vivo* cortical modules, we could expect other features. For example, during phase (i) the increasing *r*_L6_ brought to a shallower Down valley and thus a smaller restoring force, which in turn made the system wandering slower and with wider fluctuations (see the rise of variability across single gray traces). As result, the power spectral density (PSD) *P*(ω) of *r*_L5_ had to display a relatively large increase of power at low-ω band (LFB) compared to the power increase at high-ω band (HFB) proportional to the slight rise of *r*_L5_ (**Fig. 5e**). A similar spectral modulation was previously found in intracortical recordings from behaving monkeys^44^, where a spectral modulation index SMI = (Δ*R*_HFB_ − Δ*R*_LFB_)/(Δ*R*_HFB_ + Δ*R*_LFB_) resulted to be negative (SMI < 0) when an attractor state was destabilized as in phase (i) of the hysteresis loop. Here, Δ*R*_LFB_ (Δ*R*_HFB_) was the average ratio of *P*(ω) in the frequency band [50,250] Hz ([250,1100] Hz) between the beginning and the end of phase (i), i.e. bottom and up relative PSD in (**Fig. 5e**), respectively (Online Methods). In phase (iii), when L5 module was in Up state and *r*_L6_ decreased, the opposite trend should be expected (**Fig. 5d**): increasing *r*_L5_ had to correspond to a narrowing Up valley, and hence to a more stable Up state, yielding to SMI > 0.

Another important feature that we introduced in the L5 minimal model was an activity-dependent fatigue mechanism capable to reproduce the spike frequency adaptation systematically observed after Up state onset (**Fig. 1f**). Relying on a mean-field approximation^45–48^, a fatigue variable *c* was used to modulate an inhibitory feed-back to L5 module, which increased during sustained firing of Up states (**Fig. 5f**, Online Methods). Such fatigue variable aimed to account for changes in ionic concentrations (e.g. calcium, sodium) modulating afterhyperpolarization potassium currents^24^ or synaptic short-term depression^49^, and followed an integrator dynamics: τ_c_ d*c*/dt = − *c* + *r*_L5_. As a result, the fatigue of the L5 module progressively accumulated during the Up states, with a corresponding rise of adaptation or functional self-inhibition. As a result, Φ shifted to the right in **Figure 5b**, and hence the same did the nullclines in **Figure 5g**, where the force field vanishes and an asymptotic *r*_L5_ was expected. In other words, due to the activity-dependent adaptation modulated by *c*, the L5 effective energy landscape changed across the Up state and the activity threshold for *r*_L6_ determining *r*_L5_ Up-to-Down transitions increased, eventually reducing the hysteresis loop area. This brought to another prediction to test: Up states with increasingly long durations would raise the deactivation threshold, resulting into a progressive reduction of the hysteresis area, as shown for the model in **Figure 5h**.

### Time- and depth-dependent excitability of V1 cortical columns

In order to test the predictions made by the model, we first explored the modulation in time of the excitability of L5 across all experiments. All Up states were sorted by duration and pooled into groups. For each group the average L5 and L6 MUAs was computed as in **Figure 4a**. In the majority of the recordings (n = 16 out of 18) we found a significant decrease of the hysteresis area for increasing Up durations (*P* < 0.001, Wilcoxon signed-rank test; for an example see **Fig. 6a**). The excitability modulation was remarkably similar to that predicted by our minimal rate model in **Figure 5h**. In particular, the ascending phase (i-ii) of the loop was almost unaffected by the Up state elongation, contrary to what occurred to the descending part of the loop (**Fig. 6a**). This further supported the hypothesis of a history-dependent modulation of the Up state deactivation threshold (**Fig. 5g**). At the population level, this is confirmed by the relationship between the average relative change of hysteresis area with respect to its mean against the different Up states (**Fig. 6b**).

**Figure 6.**
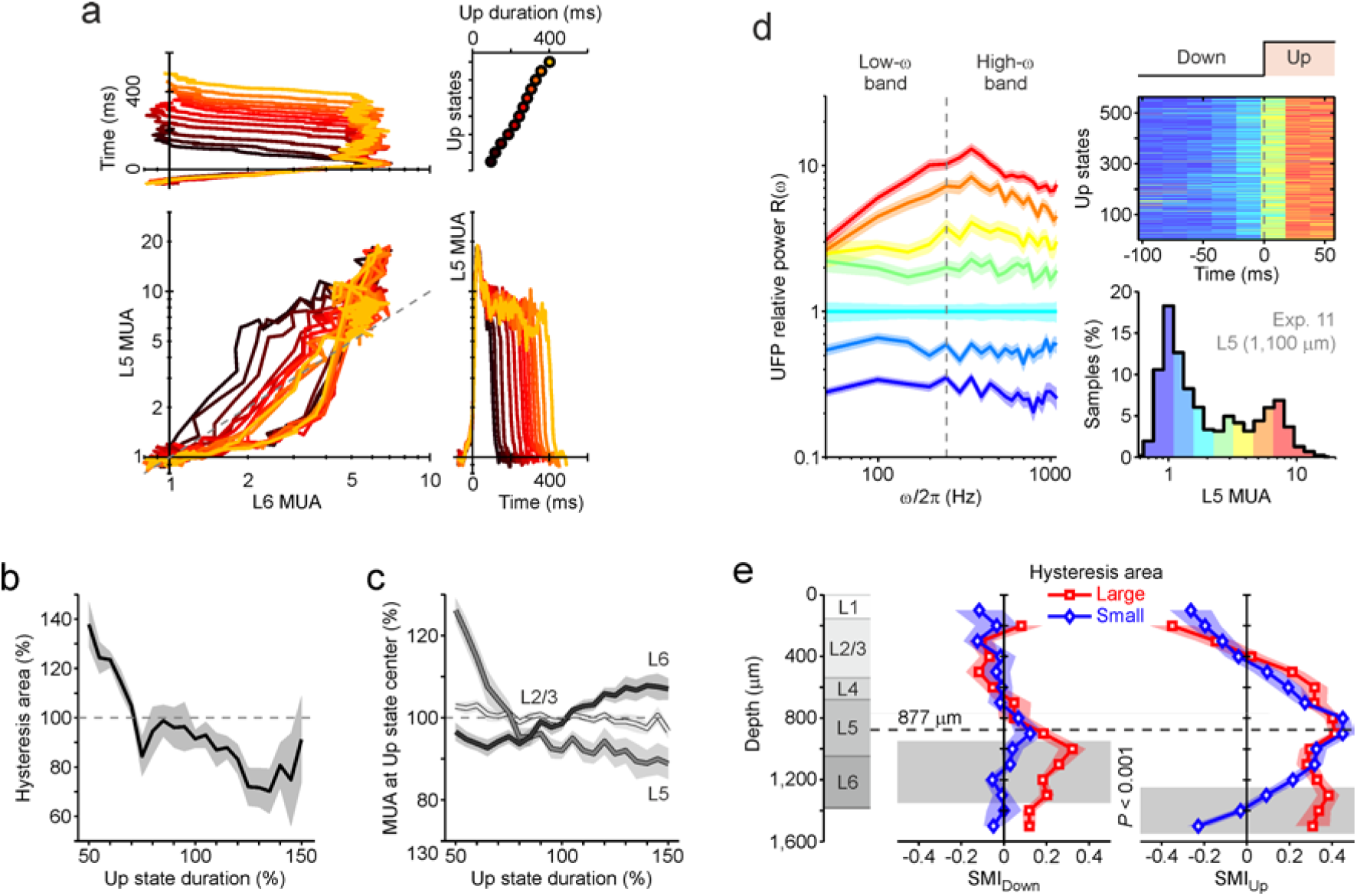
History-dependent and layer-specific cortical column excitability. (**a**) L6 and L5 MUA averages on Up state groups with different durations, computed as in **Figure 4a** for the same example recording. Up states were sorted by duration and pooled in groups of 100 states. Different groups were shifted with steps of 40 Up states. Increasing Up durations were color coded from dark brown to orange, as in **Figure 5h**. (**b**) Relative L5-L6 hysteresis area at different Up durations averaged across experiments (n = 18). Percentages of hysteresis area and Up duration were with respect to averages across all Up groups in the same experiment. (**c**) MUA measured at the center of average Up states (as in **a**) *versus* Up state duration, average across experiments. MUA and durations were expressed in percentage of averages on single experiments as in **b**. MUA were from L2/3, L5 and L6. (**d**) Relative power spectra *R*(ω) of unfiltered field potentials (UFP) estimated in 20 ms time window within [-110,70] ms around L5 Up state onsets. The same experiment as in **a**. MUAs in these time windows (top-right) were pooled in 7 groups (color coded as in bottom-right histogram). Average power spectra *P*(ω) were carried out for each group. *R*(ω) for each MUA level (left) was the ratio between corresponding *P*(ω) and the one at a reference level (third MUA level). (**e**) Spectral modulation index SMI across cortical depth, averaged across experiments with largest (red, n = 6) and smallest (blue, n = 6) hysteresis areas for short Up states as in **Figure 4**. SMI_Down_, SMI for the first three (bluish in **d**) MUA levels during Down state. SMI_Up_, SMI for the last three (yellow-reddish in **d**) MUA levels during Up state. Gray shadings, where blue and red averages were significantly different (*P* < 0.001). Reference depths for layers and dashed line as in **Figure 1h**. Colored shadings, s.e.m.

We also looked at the relationship between Up state duration and firing rate across layers (**Fig. 6c**). Only L5, the layer with higher firing rates and nonlinear dynamics, displayed adaptation of the MUA in relation with the Up state duration. This feature was not observed in L2/3 and L6. This diversity in the firing rate dynamics across cortical layers was further investigated by measuring the power spectrum modulation of the firing activity around Down-to-Up transition, in order to characterize the stability and attractor features of neuronal pools in the cortical column. We estimated the power spectra *P*(ω) of the unfiltered field potentials (UFP) in small non-overlapping time windows of 20 ms around the detected MUA upward transitions (**Fig. 6d**, top-right). We pooled such time windows in 7 groups depending on their MUA (**Fig. 6d**, bottom-right), eventually carrying out average spectra in each of them. Finally, for each activity level we computed the relative spectra *R*(ω) using as reference *P*(ω) from the third MUA group (**Fig. 6d**, left). As argued in ref. ^37^, we interpreted such *R*(ω) as the relative power spectra of the firing activity at the different stages of SO in order to test our model predictions. Specifically, the first three levels (blue range bins) around the Down state low-frequency peak of the MUA histogram, corresponded to phase (i) of hysteresis loop (**Fig. 5c i**, to compare with **Fig. 5e**). The last three MUA levels (yellow-red bins) around Up state peak represented phase (iii) of the loop (**Fig. 5c iii**, to compare with **Fig. 5d**). From these activity-dependent *R*(ω) we worked out the SMI defined above, for both the Up (SMI_Up_) and Down (SMI_Down_) activity levels and for each available electrode at different depths, eventually averaging out across two groups of experiments: the first (last) one-third of the experiments with the smallest (largest) hysteresis areas (**Fig. 6e**).

As predicted, the most stable (deep valleys in the effective energy landscape) neuronal assemblies during Up states were found in L5, where the hysteresis loop occurred and the highest MUAs were recorded (dashed line). During Down states, the power spectra were almost not modulated (SMI ≈ 0) from L2/3 to upper L5. This was incompatible with a scenario where the Down-to-Up transition would be primed by a progressive destabilization of L5 Down state as in **Figure 5c i**. Rather, an external input brought L5 assemblies to cross the energy barrier and fall into the Up state. On the other end, SMI_Up_ in supragranular layers became negative suggesting an increase of slow fluctuations in the firing rates, and thus with weaker restoring forces during Up states. Interestingly, the main differences between the two group of experiments with small and large hysteresis area were found in deep layers (mainly L6). Those with large loops displayed significantly (*P* < 0.001, Wilcoxon rank-sum test) more stable Up and Down states (SMI > 0). In particular, L6 spectral modulation during Up states was comparable to the one found in L5.

### Cortical origin of Up states and role of L6 and thalamus

SO are the local expression of a collective thalamocortical network where Up states propagate as slow waves across cortical areas^7,11,13,15^. Afferent projections to V1 cortical columns are from striate cortex itself, from other cortices at higher hierarchical levels^29,41^, and from LGN, the early thalamic station processing visual stimuli. To disentangle the roles of cortex and thalamus, we inactivated the LGN ipsilateral to the previously recorded V1 (n = 7) (Online Methods). The local injection of TTX in the LGN caused a total block of visually-evoked cortical responses (not shown).

The spatiotemporal patterns of MUA (**Fig. 7a**) and synaptic activity (**Fig. 7B**) characterizing Up state onset across cortical layers were not affected by LGN inactivation. In particular, under “LGN off” condition, the L6 MUA pre-activation systematically occurred in all experiments, displaying the same tight correlation with the onset time of the CSD sink in L6 found under control condition (**Fig. 2d**). These observations suggested that the synaptic input to L6 modules, that primes Up state activation in the column was of cortical origin, further supporting the hypothesis that the thalamus had a vicarious role in generating SO in sensorial areas^13,15^. Besides, we found no significant changes in the dynamic properties of L5 modules (**Fig. 7c**), like the hysteresis area (P = 1, Wilcoxon rank-sum test) and hysteresis modulation by Up duration before and after the LGN blockade. This evidence supports the hypothesis of a local cortical origin of the attractor dynamics expressed by L5 modules, as found in *in vitro* preparations^16,39,50^.

**Figure 7.**
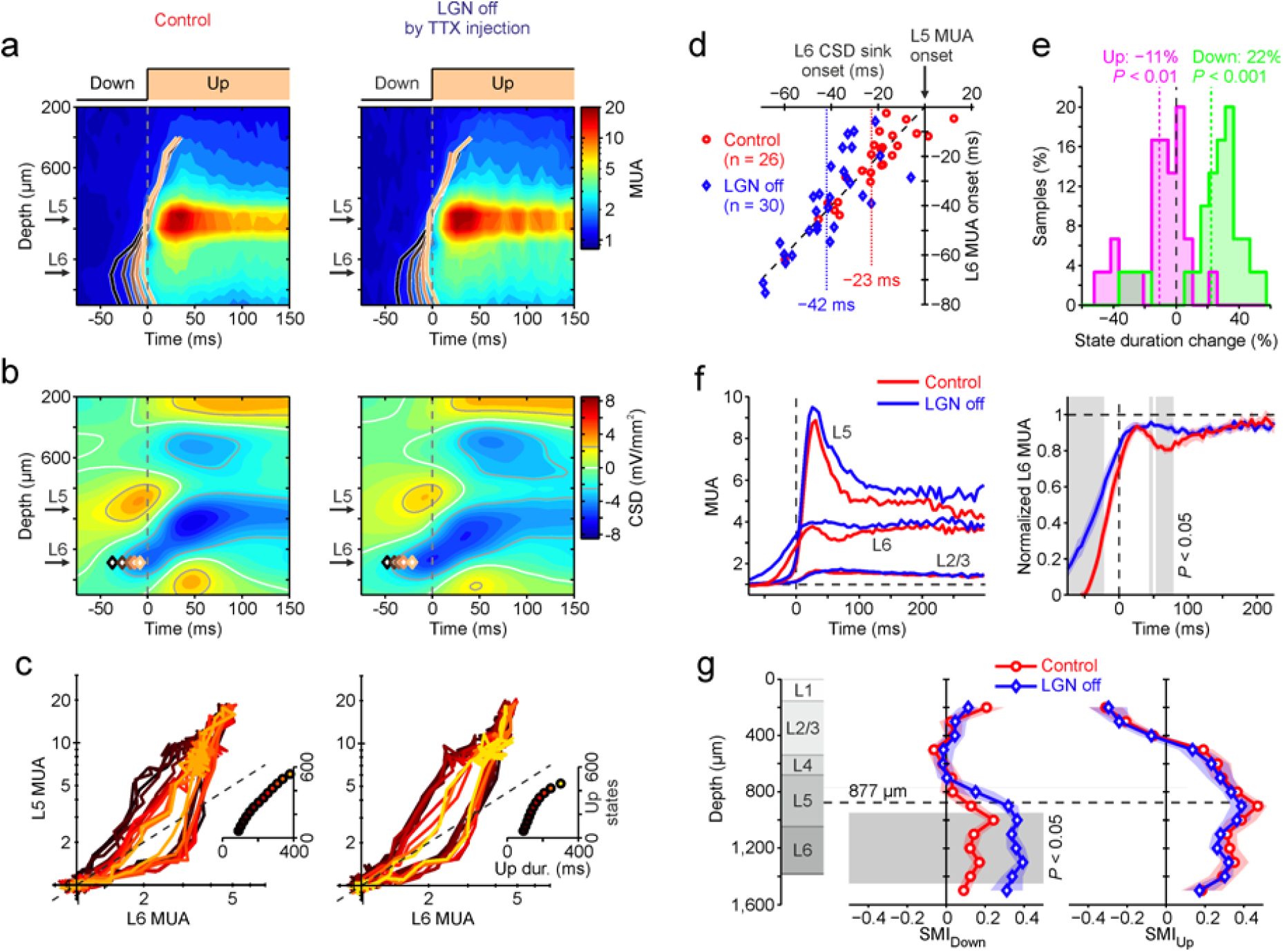
LGN inactivation dampens the excitability of deep cortical layers. (**a**) Average MUA around L5 Up state onset time under control (left) and LGN off (right) conditions for an example recording. Brown curves, average MUA onset times over grouped upward transitions as in **Figure 2c**. (**b**) Average CSD centered around Up onset times as in **a**. Brown diamonds, onset times of CSD sink in L6 for the same groups of upward transitions in **a**. (**c**) Hysteresis loop in the plane L6-L5 MUA at different Up state durations (insets) as in **Figure 6a**, for the same example recording. (**d**) L6 CSD sink and MUA onset times across all upward transitions groups from all experiments (n = 7) with both control (red) and LGN off (blue) conditions. Onset times with respect to Up state activation time in L5. Dashed line, where X = Y. Colored dotted lines, average onset times in the two data sets. (**e**) Histograms of relative change in Up (magenta) and Down (green) state duration before and after TTX injection for the same groups of upward transitions in **d**. Reference durations, those from control condition. Dotted lines, averages. (**f**) Average MUA around Up state onset for layers 2/3, 5 and 6 in both experimental conditions across all experiments (n = 7). Right, average L6 MUA normalized to the asymptotic activity levels from each experiment. Gray strips, significant differences between conditions (*P* < 0.05, Wilcoxon rank-sum test). (**g**) Average SMI across experiments (n = 7) during Up (right) and Down (left) states as in **Figure 6e**. Colors as in **d** and **f**.

Even when the LGN blockade did not alter Up states initiation, it did have some significant effects in the columnar dynamics. We first found differences in the activation of deep cortical modules. L6 CSD sink and L6 MUA onset occurred almost simultaneously, however after the LGN blockade, the delay of L5 MUA onset almost doubled from −23 ms under control condition, to −42 ms (**Fig. 7d**; *P* < 0.001, Wilcoxon rank-sum test).

Up and Down state durations significantly changed (**Fig. 7e**), such that after TTX injection Up states were shortened by −11%, while Down states were elongated by +22% (*P* < 0.01 and *P* < 0.001 respectively, Wilcoxon rank-sum test). Both these results pointed out to a possible decrease of cortical excitability. We tested such hypothesis by inspecting the average time course of MUA in different layers around the Up state onset in L5 (**Fig. 7f**). No significant differences were found in L5 and L2/3 MUA. We found though a significant increase of L6 MUA at specific time intervals as a consequence of silencing the LGN (**Fig. 7f**, right). Specifically, L6 MUA displayed an earlier activation preceding Up state onset in L5 (**Fig. 7d**), with a slower slope than in control condition. Furthermore, an indent in the L6 MUA observed in control condition after about 60 ms from Up state onset, was lost in the blocked LGN (**Fig. 7f**; *P* < 0.05, Wilcoxon rank-sum test).

By looking at the SMIs of cortical modules before and after TTX injection into LGN (**Fig. 7g**), we found that SMI_Down_ displayed a significant increase for LGN off (P < 0.05, Wilcoxon rank-sum test) in the deeper part of L5 and in L6, while SMI_Up_ remained almost unchanged. The absence of synaptic inputs from LGN therefore yielded to a more stable Down state in deeper cortical layers and hence less prone to elicit Up states of the whole cortical column. This is coherent with the longer Down states and the slower initiation of Up states and with the apparent decreased excitability.

## DISCUSSION

Examining at high spatiotemporal resolution the spiking and synaptic activity of neuronal assemblies across a cortical column we found a hierarchy of functional loops, mirroring a multiscale organization of slow wave activity. Recurrent synaptic connectivity in L5 assemblies implemented the smallest of these nested loops (**Fig. 8a**). When the cortical network is isolated in slices maintained *in vitro*, such local connectivity enables a collective attractor dynamics with two alternatively visited metastable states of persistent reverberating activity (Up) and quiescence (Down) ^15,38,40^. We found *in vivo* a similar nonlinear dynamics, with a main difference: Down states in L5 were stable attractors, and Down-to-Up transitions were primed only by a large enough input, which here we found to originate in L6. As expected for these kind of nonlinear systems, L5 assemblies gave rise to hysteresis loops, whose areas reflected their degree of excitability and their capability to have history-dependent responses (**Fig. 8a**, bottom). Adaptation mechanisms damped L5 spiking frequency down during Up states, yielding to a shrinkage of hysteresis loops, i.e. to a reduced excitability, and eventually to the loss of bistability for long enough Up periods.

**Figure 8.**
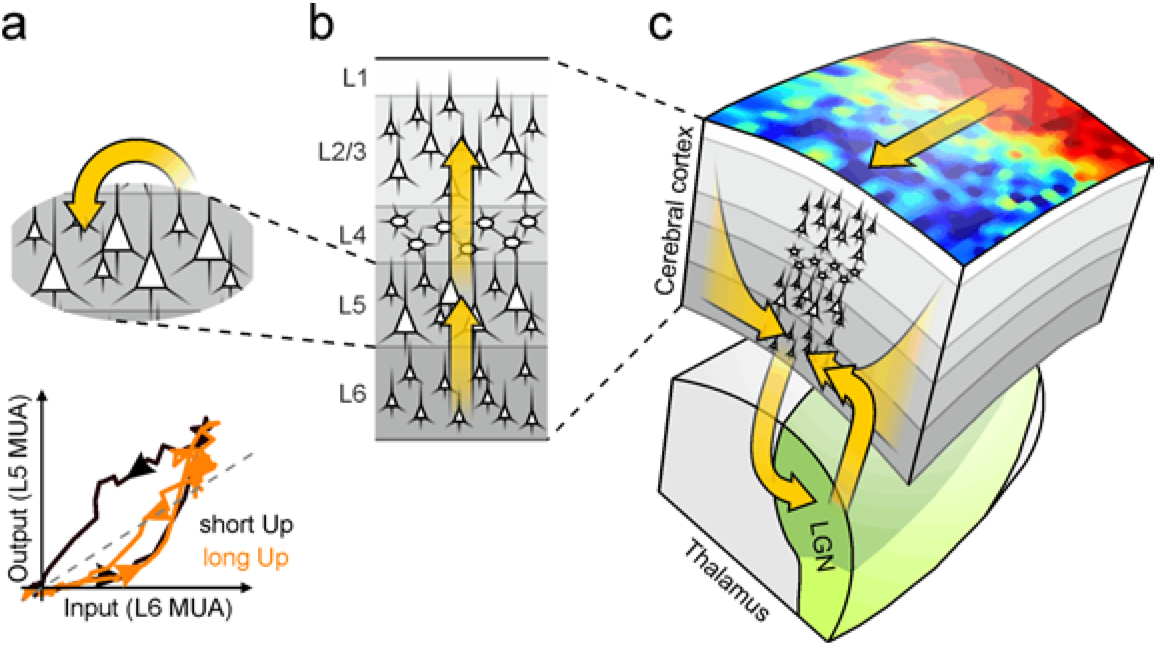
Multiscale organization of slow oscillations. (**a**) At the lowest spatial scale are the single L5 neuronal assemblies, the engines of SO, where two attractor Up and Down activity states coexist thanks both to their intrinsic cell properties, and to intense local synaptic reverberation. Such collective nonlinear dynamics underlie the measured hysteresis loop (bottom, from **Fig. 7c** left), differently expressed during long and short Up states because of spike frequency adaptation or other activity-dependent fatigue mechanisms. (**b**) At the larger spatial scale of cortical columns, activity onset flows upward from L6 to more superficial layers determining a reproducible chain of orderly activations. (**c**) At the highest macroscopic scale are located the input sources eliciting and further modulating SO at the columnar level. Synaptic inputs priming L6 activations are of cortical origin, likely originated by nearby columns involved in the propagation of Up state wavefronts across the cortex. Columnar activity is fed back to the connected subcortical structures (in this case LGN), which in turn affects the activity of L6 assemblies and modulates their excitability. Here, a traveling Up state wavefront is represented as a distribution of color coded MUA on top of the cortical surface. In all panels yellow arrows represent activity flow, likely transmitted through direct synaptic connections.

In agreement with previous reports^10,16,20,22,39^, we also found a propagation of L5 activation towards more superficial, L2/3 layers (**Fig. 8b**). The novelty here was that this stereotyped ascending pathway of Up state onsets across lamina, systematically started more in depth: from L6 with an ordered upward spreading towards the cortical surface. As the whole column was silent before its activation, only the upward branch of the bidirectional connectivity^26,30,51,52^ between L6 and L5 was involved in the chain reaction behind Down-to-Up transition, which in turn had to be primed by the excitatory synaptic input provided to L6 from other areas. Both cortico-cortical synaptic pathways^30,52– 55^, and thalamo-cortical projections to deep layers^26,28,56^ could contribute, highlighting that SO at the columnar level were only a mesoscopic component of other partially overlapped functional loops at more macroscopic spatial scales. The loop we that here contributed to unravel was cortico-cortical, as the inactivation of the ipsilateral LGN by TTX injection did not change the stereotyped spatiotemporal pattern of multi-unit and synaptic activity across the cortical column (**Fig. 7a-c**). Hence, the input to L6 had to originate from distant and nearby cortical regions^32,57,58^ (**Fig. 8c**), likely from those columns contributing to the propagation across the cortex of the Up state wavefronts underlying SO^7,11,13,59^. Although the prominence of this cortico-cortical loop ruled out a major role of the first-order sensory thalamus in the generation of such slow rhythms in visual cortex^12,13,15^, we found LGN to influence deep layers both in stabilizing Down states (**Fig. 7g**) and in affecting L6 MUA time course around Down-to-Up transitions (**Fig. 7f**). This evidence recasts the cortico-thalamo-cortical loop as also a modulator of SO in sensorial areas, strengthening the role of L6 as a fundamental hub (**Fig. 8c**) of neuronal activity produced at macroscopic scale. A role expressed not only in response to sensory stimulation^27,28^ but also under self-sustained or spontaneous state like slow wave activity.

Several studies investigated the depth profile of spontaneous columnar activation during slow oscillations^10,16,20,21,59,60^. In cortical slices displaying Up/Down oscillations, active states originated in L5 rapidly spreading upward and downward along a cortical column^16,22,39^. The same layer has a leading role in the horizontal propagation of this activation along the whole cortical slice^22,39^. Similarly, *in vivo* recordings from anaesthetized and naturally sleeping animals identified the first onset of Up states in deep/infragranular layers^10,38^, with an almost simultaneous MUA onset in L6 and L5^10,20^. By looking in detail into the activation of deep layers, here we measured a systematic pre-activation (up to a maximum of 45 ms) of L6 cortical modules preceding the activity onset in L5 where maximum firing rates were always found (**Fig. 2e**). Such deepest origin of the Up states compared to that found in *in vitro* cortical networks could be due to the loss of cortico-cortical connections in the slices plus the lack of thalamocortical inputs. Indeed, the cutting of slices affects mainly the long-range excitatory synaptic connectivity^61^, yielding to a strong reduction of the cortico-cortical input that we found to underlie L6 pre-activation *in situ*. Hence, lacking cortical slices long range intracortical connections, the generation of Up states is driven by the most excitable layer, L5, and its local connectivity driving the spontaneous Down-to-Up transitions^39,45^. Such spontaneous waves can then propagate to the nearby columns by priming other L5 activations via intact short-range intracortical connectivity^22,39^.

The role of L6 in the initiation of Up states has not been explicitly reported in previous studies *in vivo*. The maximum firing rate of the spiking activity has been reported to occur in L5 by other authors and by ourselves and in both cats, mice and rats^10,20,38^. In ref. ^20^ a second activity bump at lower spiking frequency was apparent in L6 in recordings from primary auditory cortex of urethane-anesthetized rats. As in our experiments, the laminar profile of CSD during Up states displayed an extended current sink in midlayers, confined by two narrow sources in superficial and deep layers in both cats and rats^10,20,60^. Thus, the difference in the depth of Up state origin likely lays in the adopted detection method and the sorting of state transitions, which allowed us to improve the temporal resolution in probing synaptic and spiking activity changes. For instance, the laminar activation profile of MUA in ref. ^20^ relied on the activity peak (median spike time) within each Up state at different depths. In ref.^10^ the LFP deflection at the deepest electrode was used as onset time similarly to ref. ^60^, where the negative peak in deep LFP was the reference time. All these measures might introduce time shifts with respect to more local estimates like those relying on MUA, as the approach adopted here, such that averages across Up states might mask the early activation in L6.

Our findings of *in vivo* L5 cortical modules expressing nonlinear dynamics where Up and Down states were metastable attractors, confirmed and extended the evidence resulting from *in vitro* studies^16,50^. The need of a strong enough excitatory volley to elicit a Down-to-Up transition underlying the initial branch of the hysteresis loops (c.f. **Fig. 4a, 6a** and **7c**), is in full agreement with the all-or-none response to optogenetic stimulation of L5 neurons found in anesthetized mice^13,21^. Under similar stimulation conditions, *in vivo* L2/3 cell assemblies were not capable to elicit neither locally nor in the whole cortical column an active state^21^. This scenario is compatible with an almost linear input-output response function of the cortical modules at this depth, likely underlying the absence of LFP spectral modulations in both Up and Down states (SMI ≈ 0, **Fig. 6e**). On the other hand, at L5 depth SMI_Up_ was markedly positive, further denoting the onset of a local attractor dynamics in the cortical modules during active states. This fingerprint left by the local network activity was the same as the one previously found in premotor cortical modules recorded from macaque monkeys performing reaching tasks^44^, although referring to a different brain state. Such widespread evidence further corroborates the hypothesis that flip-flop neuronal assemblies can be found also across the intact neocortex, a kind of universal computational unit to use in composing macroscopic spatiotemporal patterns of activity like traveling waves during deep sleep and anesthesia or distributed activity onsets related for instance to a behavioral output.

The metastability of Up states in L5 modules was found to be contingent on the duration of persistent spike firing (**Fig. 6a-c**). This history-dependent modulation of the network excitability was fully captured by our minimal rate model, which effectively integrated adaptive mechanisms previously introduced in more detailed network models of SO^23–25^. The gain modulation underlying such Up state destabilization, was thought to cause the sharp Up-to-Down transition, but here we found that spike frequency adaptation in L5 reached a plateau in those Up states with duration longer than the population average, i.e. roughly after 150 ms from columnar activation (**Fig. 6c** and **7f**). Beyond this time lag, hysteresis loops were widely shrunk (**Fig. 6b**) and high-firing in L5 was likely sustained only thanks to the input coming from the persistent spike firing of the bottom L6 module. This hypothesis was further supported by the evidence that in long Up states, Up-to-Down transitions in L6 occurred simultaneously or later than in L5 (**Fig. 3g**). Intriguingly, the bistability of L5 modules was also heterogeneously expressed across experiments: the maximal area of the hysteresis loop was different from column to column (**Fig. 4b**). As if a wide spectrum of columnar modules with different degrees of nonlinear input-output properties would be available, and which in turn could be further modulated in a history-dependent manner.

The inactivation of LGN did not affect the spatiotemporal profile of columnar activation (**Fig. 7a-c**), further confirming the cortical origin of *in vivo* Up state in sensory cortex as in refs. ^13,15^. We found, however, a significant albeit small (11%) reduction of Up duration following LGN inactivation widely counterbalanced by an elongation of the Down states (**Fig. 7e**). A change similar to what found in ref. ^62^, although a more dramatic reduction of the SO frequency was measured there. To this picture, our findings add details about the possible mechanistic underpinnings of such modulatory role of the first-order thalamus^63^. During LGN inactivation, the increased unbalance between active and silent state durations correlated with an increase of the stability of the Down state in deep layers (mostly L6, c.f. **Fig. 7g**), eventually reducing the excitability of the bistable L5 modules, as previously found in cortical slices^45^. and the intact brain^48^. In other words, Down-to-Up transitions have a larger chance to occur when thalamus is unblocked. Thus, the resting thalamic input lowers the threshold for the cortical columns to respond intracortical synaptic inputs involved in the Up state wavefront propagation. Notably, this gain modulation is absent during active states, this suggests that the mechanistic origin is different from the one operated by L6 in visually evoked activity^27^. Nevertheless, our results support the hypothesis that even under slow wave activity, L6 acts as a hub where convergent multimodal information from sensory thalamus^26,56^ and top-down cortical projections^30,52–55^ at the macroscopic scale can be integrated, filtered and amplified at a meso/micro scale by the bistable units in L5. This by exploiting a rather versatile columnar circuitry capable to engage parallel computational pathways determined by the cortical and/or thalamic source of the input.

## ACKNOWLEDGMENTS

This research has received funding from the European Union’s Horizon 2020 Framework Programme for Research and Innovation under the Specific Grant Agreement No. 785907 and No. 945539 (Human Brain Project SGA2 and SGA3, respectively) and from the EU FP7 FET CORTICONIC contract 600806 to MVS-V and MM; and from the Spanish Ministry of Science and Innovation (BFU2017-85048-R) to MVS-V. MM would like to thank J. Braun and M. Ruiz-Mejias for stimulating discussions.

## ONLINE METHODS

### In vivo extracellular recordings

18 adult male Wistar rats weighting 229 ± 56 g (mean ± s.d.) were anesthetized via intraperitoneal injection of ketamine (120 mg/kg) and medetomidine (0.5 mg/kg). Atropine (0.05 mg/kg) was injected subcutaneously to prevent respiratory secretions. Rectal temperature was maintained at 37°C. Two craniotomies were performed to access the primary visual (V1M) cortex (7.3 mm AP, 3.5 mm ML) and the lateral geniculate nucleus (LGN) of the thalamus (4.3 mm AP, 3.6 mm ML) of the right hemisphere^51^. All experiments were supervised and approved by the local Ethics Committee and were carried out in accordance with the present laws of animal care, EU guidelines on protection of vertebrates used for experimentation (Strasbourg 3/18/1986) and the local law of animal care established by the Generalitat of Catalonia (Decree 214/97, 20 July). Recordings of cortical activity under anesthesia were obtained with a 16-channel silicone probe (1 shank with 16 linearly spaced sites at 100 µm increments with impedances of 0.6-1 MΩ at 1 kHz (NeuroNexus Technologies, Ann Arbor, MI) introduced perpendicularly in V1 under visual guidance until the most superficial recording site was aligned with the cortical surface. Signals were amplified (Multi Channel Systems) and digitized at 10 kHz and acquired with a CED acquisition board and Spike 2 software (Cambridge Electronic Design, UK). An tungsten electrode with impedance of 1-2MΩ was lowered (4.1 mm) through the LGN craniotomy to simultaneously record thalamic activity. To evoke sensory responses in V1, a light-emitting diode (LED) was placed in front of the contralateral eye and a 1 ms-flash was automatically delivered every 5 seconds for 60 times. In a subset of 7 rats, the selective blocker of Na^+^ channel conductance tetrodotoxine (TTX) was infused into the LGN at a concentration of 10 ng/µl. The infusions were delivered through a glass electrode lowered adjacent to the tungsten electrode using an infusion pump and were repeated till the visual stimulation did no longer evoke a cortical response. Recordings under LGN inactivation were always performed within 4 hours from the beginning of anesthesia induction.

### Head-fixed recordings for anatomy and electrophysiology

Four adult male Wistar rats were included in these experiments. An initial surgery was performed where animals were anaesthetized by inhalation of isoflurane (4% induction, 1.5% maintenance) and placed in a stereotaxic apparatus (David Kopft Instruments). After sectioning the scalp, a custom head post was attached to the skull by means of anchoring screws and acrylic dental cement. After surgery, animals received a daily dose of buprenorphine (0.05 mg/kg) for two days for analgesia and enrofloxacine (10 mg/kg) for five days to avoid infection. After a recovery period of at least one week, handling and head-fixation training began. Training was performed for 5-10 sessions during which the duration of head restrain was gradually extended from 10 min to 1h. At the end of the training period animals spontaneously fall asleep during each session. On the day of the recording, a craniotomy and durotomy under isoflurante anesthesia (same protocol as above) were performed over the target area (V1M, 7.3 mm posterior to bregma and 3.5 mm lateral from midline). Recording began after a recovery period of at least 1h by means of 16-channel silicone probes (1 shank with 16 linearly spaced sites at 100 µm, Atlas Neuroengineering) coated in a solution of the lipophilic vital dye DiI (Invitrogen) for histological marking that were lowered perpendicular to the cortex until the last recording electrode was aligned to the cortical surface. Signals were amplified (Multi Channel Systems) and digitized at 10 kHz and acquired with a CED acquisition board and Spike 2 software (Cambridge Electronic Design, UK). All training and recording sessions were stopped if animals showed any sign of discomfort. At the end of each experiment animals were deeply anaesthetized (4% isoflurane for induction followed by sodium pentobartal 120 mg/kg, ip) and transcardially perfused with a solution of 4% paraformaldhyde in 1% PBS. Brains were then removed and kept in 4% paraformaldhyde until histological processing. Histology was performed to verify electrode location in all cases.

### Anatomy

After perfusion, the brains were removed, postfixed for 4 h at 4°C and then crioprotected by inmersion in a in 30% sucrose until they sunk. Subsequently brains were frozen in dry ice and cut in the coronal plane with a freezing sliding microtome. Serial sections (50 µm) were stained with the nuclear stain DAPI (4,6 diamino-2-phenylindol; Sigma, St. Louis, MO, EEUU) to reveal limits between layers and cytoarchitectonic areas. After staining, the sections were mounted with ProLong Gold Antifade Reagent (Invitrogen) and examined on a fluorescence microscope (Olympus BX51). Dapi and DiI fluorescences from all serial sections through the right primary somatosensory and right primary visual neocortex was photographed (Olympus DP70). Fiji software was used to combine the images recorded through the different channels in order to study the spatial relationship between the electrode tracts and the thickness of cortical layers.

### Data analysis

From each recording channel, MUA was estimated as the power change of the UFP in sliding windows of 5 ms ^10,37^. UFP power spectra in these time windows were computed from their Fourier transform and normalized by the median spectrum across the whole time series. The resulting relative power density averaged across the frequency band [0.2, 1.5] kHz was our MUA estimate, directly proportional to the firing rate of the neuronal pool surrounding the electrode tips under the hypothesis of a stereotyped single-unit waveform ^37^. Such estimate had the advantage to be unbiased by any threshold criterion typically adopted in extracellular recordings. MUA was smoothed by scaling logarithmically and averaging on sliding windows of 40 ms the whole time series. In each recording, MUA was normalized to have an histogram with a low-value peak corresponding to Down states at 1. For each animal and condition (LGN on or off), continuous UFP time series of 500 seconds long was analyzed. CSD analysis was performed on LFP, i.e. high-pass filtered UFP at 0.1 Hz, downsampled with a moving average on 5 ms windows, as for MUA. As in ^32^, firstly we smoothed LFP ϕ(z) at depth z > 0 with a Laplacian filter, ϕ_F_(z) = [ϕ(z + h) + 2 ϕ(z) + ϕ(z − h)]/4, where h = 100 μm (the inter-electrode distance) and with the uppermost and lowermost LFP duplicated at the boundaries of the array. Thus we interpolated ϕ_F_(z) with a cubic spline, in order to compute analytically CSD(z) = − ∂^2^ ϕ_F_(z)/∂z^2^. To characterize the cortical evoked response, we monitored the changes CSD centered around light onset (**Suppl. Fig. 1**). Firstly, we high-pass filtered MUA and CSD (second-order Butterwoth filter, cut-off frequency 3 Hz) at each depth to have reliable detection of the stimulus response time within the probed column. We detected a response onset when filtered MUA averaged across electrodes crossed a threhold value (4 s.d. of the MUA in the range [-30,10] ms around light onset). Average CSD in the 10 ms following the response time showed a prominent current sink representing stimulus-related synaptic input from LGN ^33^, and hence marking L4 depth in the 15 experiments (out of 18), with succesful photic stimulation protocol.

Up and Down state transitions were detected as the crossing times by log(MUA) of a threshold value ^10^. Logarithmic scaling allowed to have in all recordings bimodal histograms fitted by a superposition of two Gaussian distributions corresponding to the Up and Down states, respectively. The threshold value was set to 60% of the gap between the peaks of the fitted distributions. State durations shorter than 80 ms (2 times the smoothing window) was discarded (see **Supplementary Figure 2** for more details). State transitions was considered to be coincident across the cortical column if occurring within a time window of 60 ms (average across experiments, values ranged from 36 ms to 84 ms and were chosen to minimize double occurrences from the same channel). Of these columnar state transitions we discarded those having i) the majority of the channels without transitions and ii) no transition at least in one of the three channels centered around L5 depth (where maximum MUA was measured). At least 250 state transitions per experiment were collected, with an average of 438 ± 85 (mean ± s.d., n = 18). As reference time for the transitions, the average transition time across the three channels around L5 was adopted, and an array of relative time lags of state change times across the channels was computed for each transition. In those few columnar transitions with channels without detected events, the average time lag across the 5 more similar columnar transitions (in terms of Euclidian distance between arrays) was replaced. As next step, a PCA was performed on the resulting time lag matrix of each recording. The first principal component (always explaining more than 30% of the matrix variance) was used to sort the transition arrays and dividing them into 5 equally-sized groups (**Fig. 2a** and **3a**). Average MUA around state transitions in each group was fitted at each depth by a generalized logistic function σ(*t*;*a,b,c*) = (1 + e^a − b t^) ^−1/c^, in the time window [-100, 50] ms around columnar transition times, normalizing neuronal activities to their maxima. Finally, Up onsets and offsets were re-evaluated from the fitted σ(t) as the crossing time of a threshold at MUA = e^1/2^, a value two times larger than the s.d. of log(MUA) during the Down states in all the recordings. Grouped transition times of representative L5 channels were compared with those at L6 depth, i.e. where a second lower MUA peak appeared during Up states (**Fig. 2c, 3c** and **7a**). We verified that columnar spatiotemporal pattern of activity around Down-Up transitions was not affected by the specific parameters chosen to estimate MUA, like frequency range and temporal windowing, trying also to use other methods like those rectifying or thresholding high-pass filtered UFP (**Suppl. Fig. 3**). Average MUA profiles of whole Up states plotted in **Figures 4, 6** and **7**, and composing hysteresis loops, were obtained concatenating transition-triggered MUA averages and considering only the first and last half of the Up state duration from the onset and up to the offset time, respectively. Here, the Up duration was the length in time of the Up states at L5 depth, averaged across the selected transitions. In **Figures 6c** and **7f**, the MUA at L2/3 depth was in each recording the one from the nearest electrode to the depth of LFP polarity reversal.

For Down-to-Up transitions, average CSD was inspected at the columnar activation time in each Up state group and at depths below 950 mm (average depth of L5 center). Deep CSD sink had depth where a CSD minimum was found under a threshold value of −2.5 mV/mm^2^. Onset of this CSD sink corresponded to the downward crossing time of the same threshold value before Up state onset. The CSD threshold was chosen to be in absolute value at least two times the s.d. of the CSD at the Down state end (within [-150, −50] ms from the activation time) across all recording channels. The depth of the CSD sink around Up-to-Down transitions was estimated adopting the same criteria, but using as reference time t_0_ = 40 ms after columnar inactivation (Up state offset). The offset of these CSD sinks, corresponded to the upward crossing time of the CSD threshold after t_0_.

### Computational model and simulations

The minimal rate model of the L5 cortical module, received an input proportional to L6 activity *r*_L6_, represented by a sigmoidally-saturated sinusoid: *r*_L6_(*t*) = σ(sin(2π*t*/*T*); 50 θ_Up_, 50, 1). This to have Up/Down oscillations in L6 at frequency 1/*T*, as shown in **Figure 5**. The duty cycle between Up and Down states was regulated by θ_Up_ = sin(π/2 − π *T*_Up_/*T*), with *T*_Up_ the Up state duration. In **Figure 5h**, we set *T*_Up_/*T* = 0.25 and *T* ranged between 0.5 s to 2.0 s. Here, firing rates were scaled to have a maximum activity at 1. In the stochastic first-order dynamics of L5 activity *r*_L6_ [τ d*r*_L5_/dt = Φ(*r*_L5_^(N)^, *r*_L6_, *c*) − *r*_L5_], the sigmoidal input-output gain function was Φ(*r*_L5_^(N)^, *r*_L6_, *c*) = σ(*r*_L5_^(N)^, *I*_0_ − *w*_56_ *r*_L6_ + *g*_c_ *c, w*_55_, *k*), where under mean-field approximation, the state-dependent input was a linear combination of currents: the synaptic current from L6 modulated by the efficacy *w*_56_, the after-hyperpolarizing current given by *g*_c_ times the adaptation level *c*, and the background current *I*_0_. Recurrent synaptic input within L5 was modulated by the efficacy *w*_55_, determining the steepness of Φ, and hence the input amplification by the L5 module. Parameter *k* shaped the asymmetry of the attraction basins of the Up and Down metastable states. To include the finite-size fluctuations needed to have physiologically plausible activity power spectra within a metastable state (**Fig. 5d-e**), *r*_L5_^(N)^ = *r*_L5_ + *N*^−1/2^ *W*(*t*) was the instantaneous firing rate of the finite-size L5 module, with *N* the number of neurons in the pool and *W*(*t*) a Gaussian white noise with zero mean and unit variance. Finally, the fatigue level *c*(*t*) had the following integrator dynamics τ_c_ d*c*/dt = − c + *r*_L5_^(N)^. Both the stochastic differential equations for *r*_L5_ and *c* were numerically integrated resorting to the Euler method with a time step of 0.1 ms. In the illustrative example shown in **Figure 5b-e**, we set *N* = 250, τ = 14.4 ms, k = 1.02 − 0.28 *r*_L6_, *I*_0_ = 3.27, *w*_56_ = 1.31 and *w*_55_ = 5.26 + 0.97 *r*_L6_, neglecting the effect of the adaptation dynamics (*g*_c_ = 0). In the other panels of **Figure 5**, no finite-size noise was embodied (*N* → ∞), setting τ = 2 ms, k = 1.1, *I*_0_ = 3.5, *w*_56_ = 1.2 and *w*_55_ = 6, and for the adaptation dynamics *g*_c_ = 1.5 and τ_c_ = 100 ms. In order to express L5 model activity in the same units adopted for the *in vivo* MUA, we exponentially mapped the modeled activity as MUA_L6_ = exp(1.75 *r*_L6_) and MUA_L5_ = exp(3 *r*_L5_ − r_Down_), where *r*_Down_ was the average *r*_L5_ across Down state. Effective energy landscapes in **Figure 5c** were obtained integrating over *r*_L5_ the driving force Φ(*r*_L5_) − *r*_L5_. As in ref. ^37^, power spectra *P*(ω) of *r*_L5_^(N)^ had an analytic expression when the module activity stochastically wandered across the energy valley of a metastable state: *P*(ω) ∝ *r*_L5_ (1 + ω^2^ τ^2^) /[(1 Φ′)^2^ + ω^2^ τ^2^]. Here *r*_L5_ was the asymptotic firing rate, the stable fixed point, in the infinite-size limit (*N* → ∞). The resulting spectra had a flat asymptote at high-ω [*P*(∞) ∝ *r*_L5_], while the low-ω component was modulated by the slope of Φ′ = dΦ/d*r*_L5_ [*P*(0) ∝ *r*_L5_/(1 − Φ′)^2^], and hence by the curvature of the attractor valley.

Numerical integration of model rate dynamics and off-line data analyses were both performed in MATLAB (The MathWorks).

## Supplementary Material

**Supplementary Figure 1.**
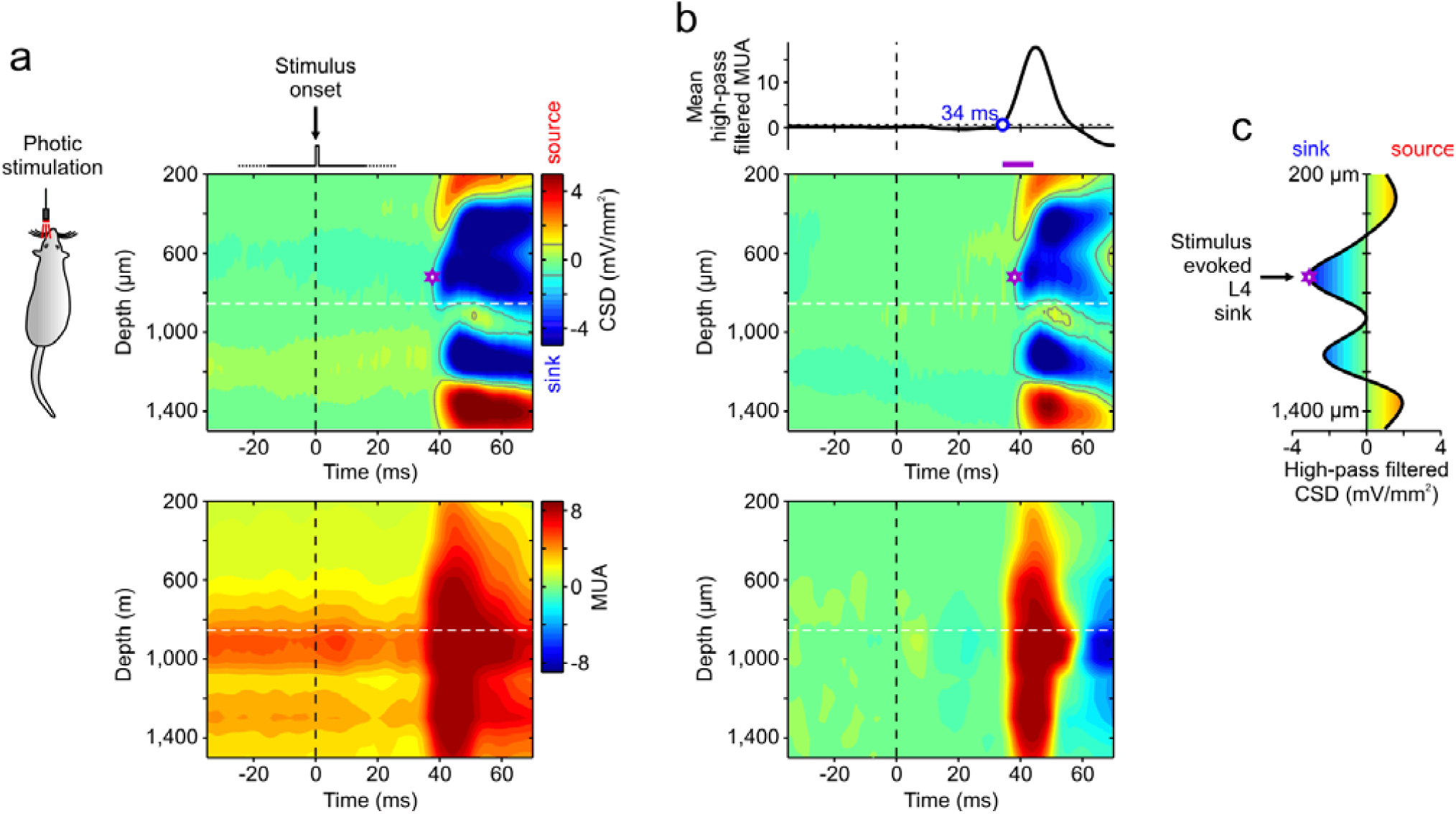
Stimulus-evoked synaptic (CSD) and spiking (MUA) activity to identify thalamic input in L4 to use as cortical depth reference. (**a**) Average unfiltered CSD (top) and MUA (bottom) in response to repeated brief photic stimulations (vertical dashed line, t = 0 ms), from the same example experiment as in **Figure 1a-d**. In each of the 15 experiments (out of 18) in which photic stimulation was successfully administered, delivering an average number of 134 ± 67 (mean ± s.d.) LED light flashes of 1 ms. A sudden change of MUA at all depths is apparent after few tens of milliseconds from stimulation time. Almost simultaneously CSD sinks and sources are measured, displaying a rather stereotypical stimulus-evoked pattern ^1,2^: an early prominent sink (above white dashed line) pointing out the thalamic synaptic input to L4, shows up followed by a more in depth source and a second deeper sink. The border (dashed white line) between L4 sink and the nearby bottom source, is just above the depth where maximum MUA spontaneously occur (L5, see bottom panel). This is in agreement with ref. ^3^ where such CSD marker was used as a rough estimate of the lower bound of L4. (**b**) To avoid any bias due to the existence of the ongoing spontaneous activity apparent even before stimulation time, both CSD and MUA were high-pass filtered (second order Butterworth filter, cut-off frequency at 3 Hz as in ref. ^1^). This filtering did not change the stimulus-evoked pattern of CSD shown in **a**. However, allowed a reliable identification of the onset time of MUA response (bottom panel), occurring after an average lag of 36.9 ± 7.1 ms (mean ± s.d., n = 15). Top panel, the onset of MUA response was the crossing time of a threshold value (4 s.d of MUA in the time range [-30,10] ms around stimulation pooling together all electrodes) by the MUA averaged across the whole column. (**c**) Average CSD in the 10 ms time window following MUA onset was used to estimate the depth of the L4 sink elicited by thalamic input in all the 15 experiments with photic stimulation.

**Supplementary Figure 2.**
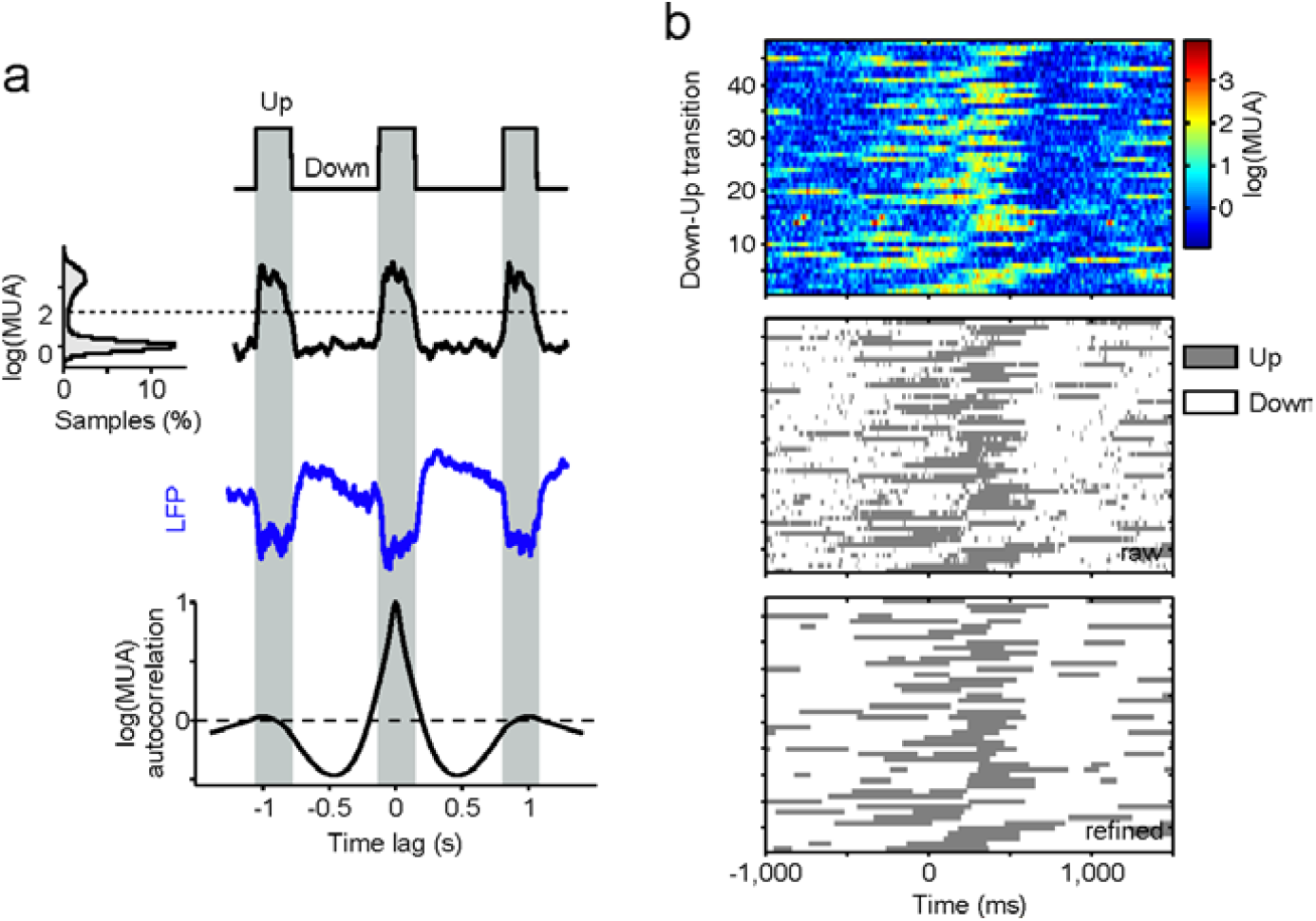
Up and Down state detection from MUA. (**a**) Sample trace of logarithmically scaled MUA (thick black) and unfiltered LFP (blue) in an example infragranular extracellular recording (same experiment as in **Fig. 1a-d**). The distribution of log(MUA) smoothed by averaging on sliding windows of 40 ms (left) is always bimodal in those recordings where MUA is maximum (L5): a feature invariant across all 18 experiments used in this work. Horizontal dotted line, MUA threshold used to detect Up-Down transitions (top trace, Up periods highlighted by vertical gray strips). This threshold is set to 60% of the gap between the two peaks corresponding to activity levels of Up and Down states. Bottom, autocorrelation of smoothed log(MUA) which in addition to the prominent peak at lag 0, displays two symmetric smaller peaks centered around the average Up/Down oscillation period. (**b**) Due to the unavoidable fluctuations of the estimated log(MUA), false positives and negatives (Up and Down state, respectively) can be detected. Top, short log(MUA) traces from an example recordings centered around arbitrary triggers selected at random times which highlight the noisy nature of the log(MUA) signal. Middle, same rasterplot in which Up and Down periods are depicted as they result from direct application of the MUA threshold to the top rasterplot. Sparse and very short false Up and Down periods are apparent together with the longer true ones. Short periods are iteratively removed if smaller than a minimum state duration, as shown in bottom rasterplot. Optimal minimum duration is the one yielding to an average Up/Down oscillation period reproducing the lag of the second peaks of log(MUA) autocorrelation (panel a, bottom). In all experiments this optimal minimum state duration resulted to be 80 ms.

**Supplementary Figure 3.**
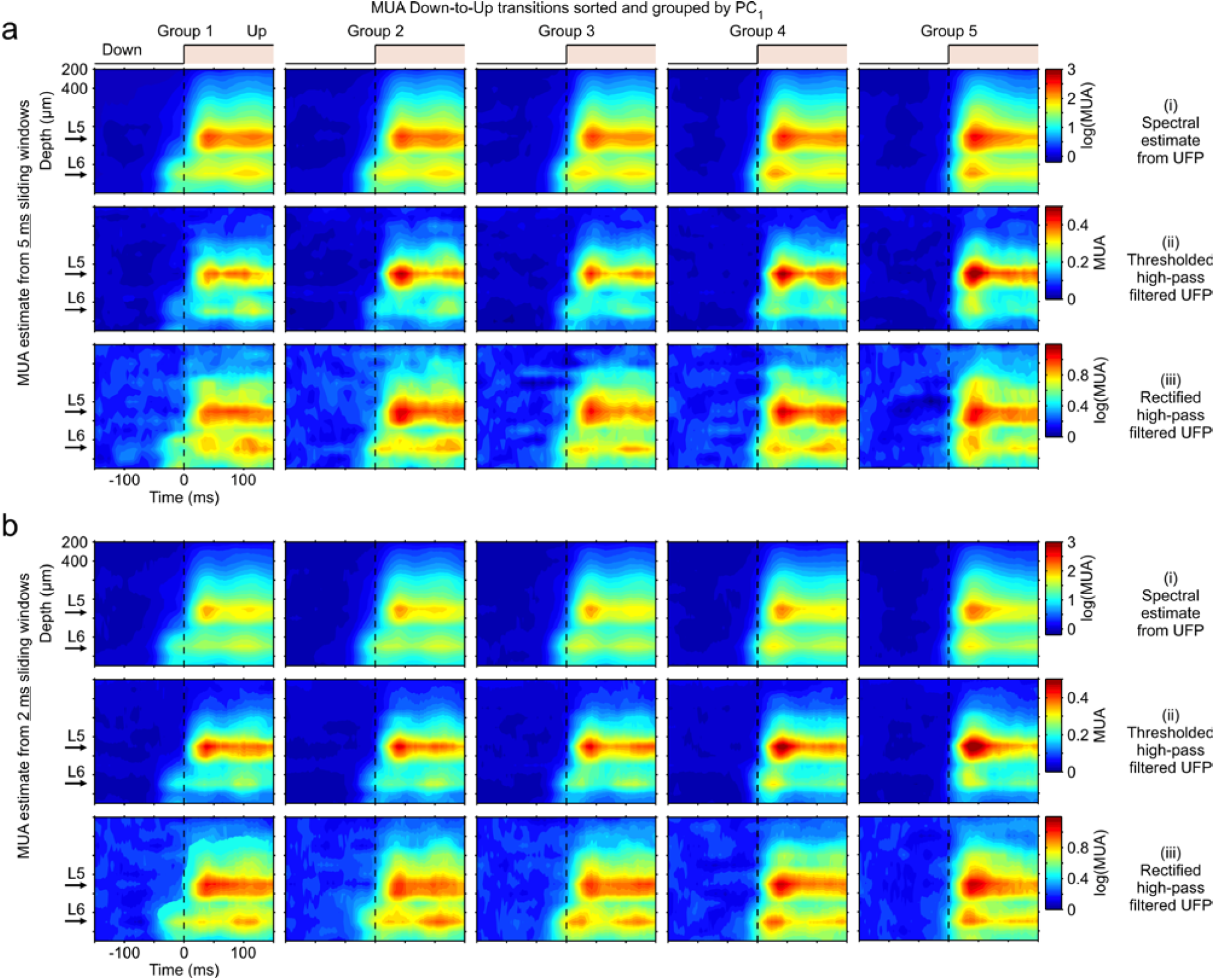
Comparison between different methods of MUA estimate. Here we aimed at investigating how much the spatiotemporal columnar activation we characterized is sensitive to the approach and parameters used to estimate MUA from extracellular recordings. To this purpose we tested and compared different MUA estimates to compute the average time course of the spiking activity around Down-to-Up transitions grouped by L6-L5 time lag, as shown in **Figure 2**. Analysis was performed on the same example recording. (**a**) MUA estimates from 5 ms sliding windows. From top to bottom: (i) MUA estimated as the power change in the high frequency range [0.2, 1.5] kHz of the UFP Fourier spectral densities, same as **Figure 2b** (see Online Methods). (ii) MUA estimated as the crossing rate of a threshold by the high-pass filtered UFP (cut-off frequency at 500 Hz). Threshold was set at 2 s.d. of the filtered UFP during Down states ^4^. (iii) MUA estimated from the rectified high-pass filtered UFP (cut-off frequency 500 Hz) ^5,6^. To remove differences in the MUA offsets resulting from different electrodes, we detrended average MUA computed within [-150, −50] ms from Down-to-Up transition. (**b**) Same as **a** but with MUA estimated from sliding windows of 2 ms. All MUA time series were smoothed computing a moving average on sliding windows of 40 ms. For high-pass filtering UFP we used a second order Butterworth filter.

**Supplementary Figure 4.**
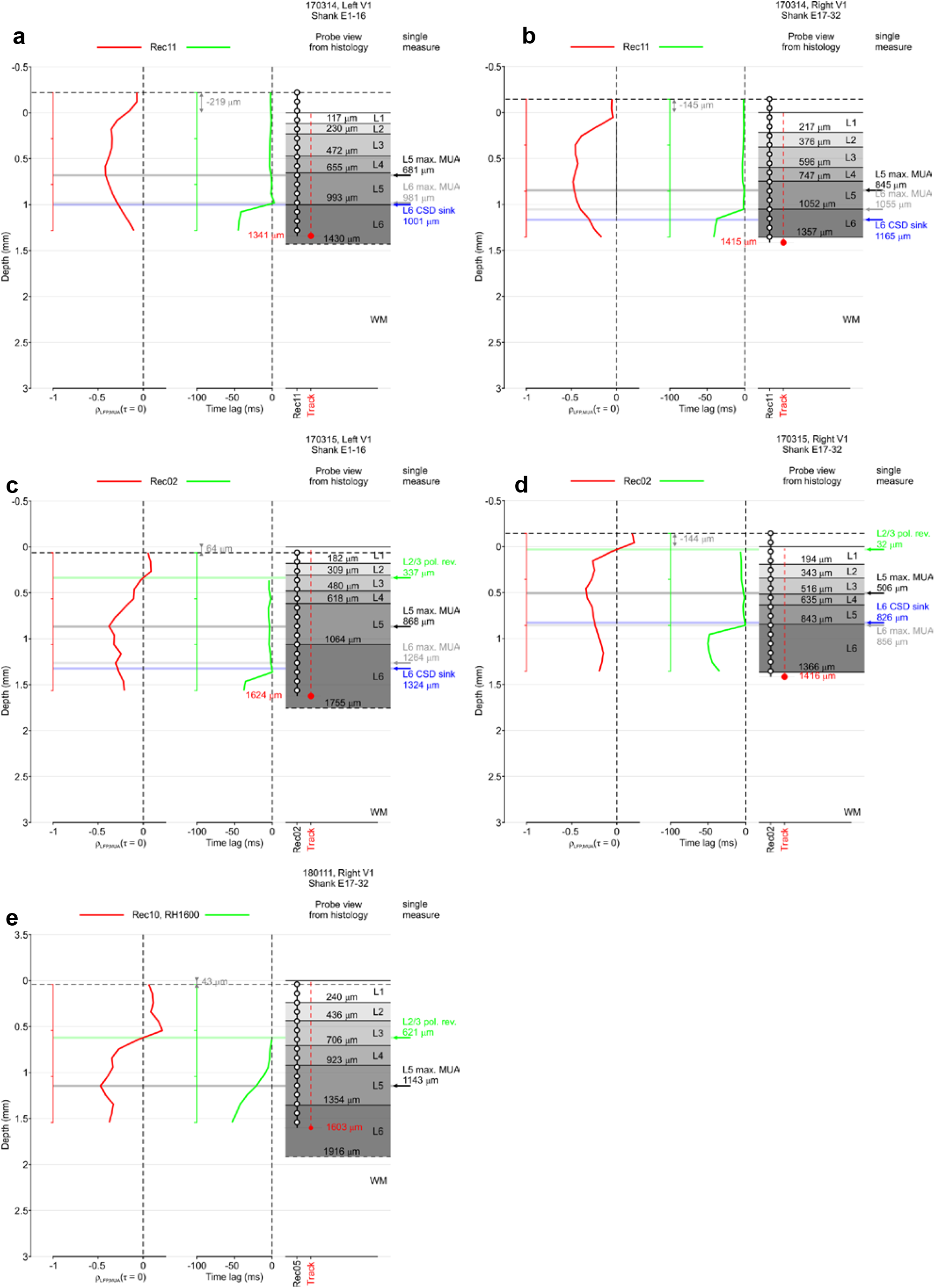
Match between electrophysiological markers obtained from MUA and CSD analysis of Down-to-Up transitions and depth distribution of cortical layers from histology. 5 **visual cortex** recordings from 3 rats anesthetized with isoflurane (the 4 topmost panels) and ketamine+medetomidine (last bottom panel). See Suppl. Fig. 5 for other details.

